# RNASTOP: A Deep Learning Framework for mRNA Chemical Stability Prediction and Optimization

**DOI:** 10.64898/2026.03.18.712573

**Authors:** Shenggeng Lin, Junwei Chen, Heqi Sun, Yufang Zhang, Wenyi Yang, Hong Tan, Dong-Qing Wei, Qinghua Jiang, Yi Xiong

## Abstract

Messenger RNA (mRNA) vaccines offer promising therapeutics for combating various diseases, yet their inherent chemical instability hampers their long-term efficacy. Although several methods have been developed to predict mRNA degradation, they exhibit limited accuracy and lack the capability for rational sequence optimization. Here, we propose RNASTOP, a novel framework integrating deep learning with heuristic search to simultaneously predict and optimize mRNA chemical stability. RNASTOP achieves a 13% accuracy improvement over the top-performing model on the Stanford OpenVaccine competition dataset and demonstrates robust generalization in predicting full-length mRNA degradation. Applied to mRNA codon optimization, RNASTOP reduces the minimum free energy of the Varicella-Zoster Virus vaccine sequence by 75.73% while maintaining high translation efficiency. Overall, RNASTOP serves as a powerful tool for predicting and optimizing mRNA chemical stability, poised to expedite the development of mRNA therapeutics. The source code of RNASTOP can be accessed at https://github.com/xlab-BioAI/RNASTOP.

## Introduction

Messenger RNA (mRNA)-based therapeutics have emerged as highly promising medical interventions [1–3], exemplified by their rapid deployment in vaccines for severe acute respiratory syndrome coronavirus 2 (SARS-CoV-2) [4–6]. However, the inherent chemical instability of mRNA poses a significant limitation to the durability of these therapeutics [7–9]. Hydrolysis within lipid nanoparticle formulations diminishes the amount of intact mRNA remaining throughout shipping and storage, while *in vivo* degradation subsequent to vaccine administration limits protein synthesis and compromises therapeutic efficacy [2, 10]. Developing mRNA therapeutics with enhanced chemical stability is therefore critical to mitigating degradation during transit and elevating overall clinical efficacy [10–12].

Codon optimization has proven to be a highly effective strategy for engineering chemically stable mRNA therapeutics [3, 9, 13–18]. Current computational optimization methods generally fall into four categories: traditional statistical approaches, heuristic search strategies, deep learning-based models, and multi-factor integration tools. Traditional approaches, such as host-preferred codon substitution, replace codons based on host frequency statistics. However, they prioritize translation efficiency while often compromising RNA chemical stability. Heuristic search-based methods, such as LinearDesign [3], utilize dynamic programming to navigate structurally stable nucleotide sequences by translating key features like folding free energy into computable scores. Deep learning-based methods are trained on massive datasets of natural sequences to capture underlying evolutionary information and intricate sequence patterns, thereby generating novel sequences not found in nature. Multi-factor integration tools, such as the commercial software OptimumGene, perform global computational optimization by integrating multiple factors, including codon preference, mRNA secondary structure, and GC content, to comprehensively optimize mRNA sequences.

Despite significant algorithmic advancements, identifying mRNA sequences with enhanced chemical stability within the vast synonymous sequence space remains a formidable challenge [19, 20]. For example, the SARS-CoV-2 spike protein antigen can be encoded by approximately 10^633^ potential mRNA sequences [3]. When utilizing heuristic search algorithms to navigate this massive sequence space, the mRNA degradation scoring function becomes crucial, as it dictates the trajectory of the optimization process [3]. A lower degradation score correlates with enhanced chemical stability [21, 22]. However, current predictive models acting as these essential scoring functions exhibit limited accuracy and lack the capability to guide rational sequence optimization [9]. Early models overly simplified the process, postulating that the cleavage probability of an mRNA linkage is strictly proportional to the unpairing probability of the 5’ nucleotide [21], neglecting the highly variable local sequence and structural contexts. While recent deep learning architectures, such as Nullrecurrent [9] and RNAdegformer [19], have advanced predictive capabilities, they still fail to bridge the gap between prediction and practical sequence design. Therefore, there is an urgent need for an integrated framework capable of both highly accurate prediction and practical codon optimization.

In this study, we introduce RNASTOP (Prediction and Optimization of mRNA chemical STability), a framework that can accurately predict mRNA degradation at both single-nucleotide and full-length levels and enhance mRNA chemical stability through codon optimization. On the Stanford OpenVaccine Kaggle (SOVK) competition dataset [9], RNASTOP achieved a 13% improvement in degradation prediction accuracy over the state-of-the-art model. Furthermore, we validated its robust generalization capability on an independent dataset of full-length mRNAs [23], where RNASTOP exhibited the highest correlation with experimental degradation values among all evaluated models. Crucially, RNASTOP provides biological interpretability by intrinsically capturing the sequence and structural motifs that govern degradation. Finally, we applied RNASTOP to optimize COVID-19 and varicella-zoster virus (VZV) mRNA vaccine sequences, successfully reducing their minimum free energy (MFE) by 20.96% and 75.73%, respectively. Collectively, these findings demonstrate the utility of RNASTOP in advancing rational vaccine design and mRNA therapeutics.

## Method

### Overview of the RNASTOP framework

RNASTOP is a deep learning-based framework designed to accurately predict mRNA degradation and enhance mRNA chemical stability via codon optimization. As illustrated in **Figure 1**, RNASTOP primarily consists of an mRNA degradation prediction model and an mRNA codon optimization module. Specifically, the predictive model integrates nucleic acid large language models (LLMs) and a structural embedding module to extract multi-source feature embeddings from mRNA sequences. These embeddings are then processed by a dual-branch feature decoupling-and-aggregating network for feature fusion and subsequent degradation prediction. Subsequently, the mRNA codon optimization module couples this predictive model with a beam search algorithm, utilizing both MFE and the codon adaptation index (CAI) as optimization criteria to concurrently improve mRNA chemical stability and translation efficiency.

**Figure 1.**
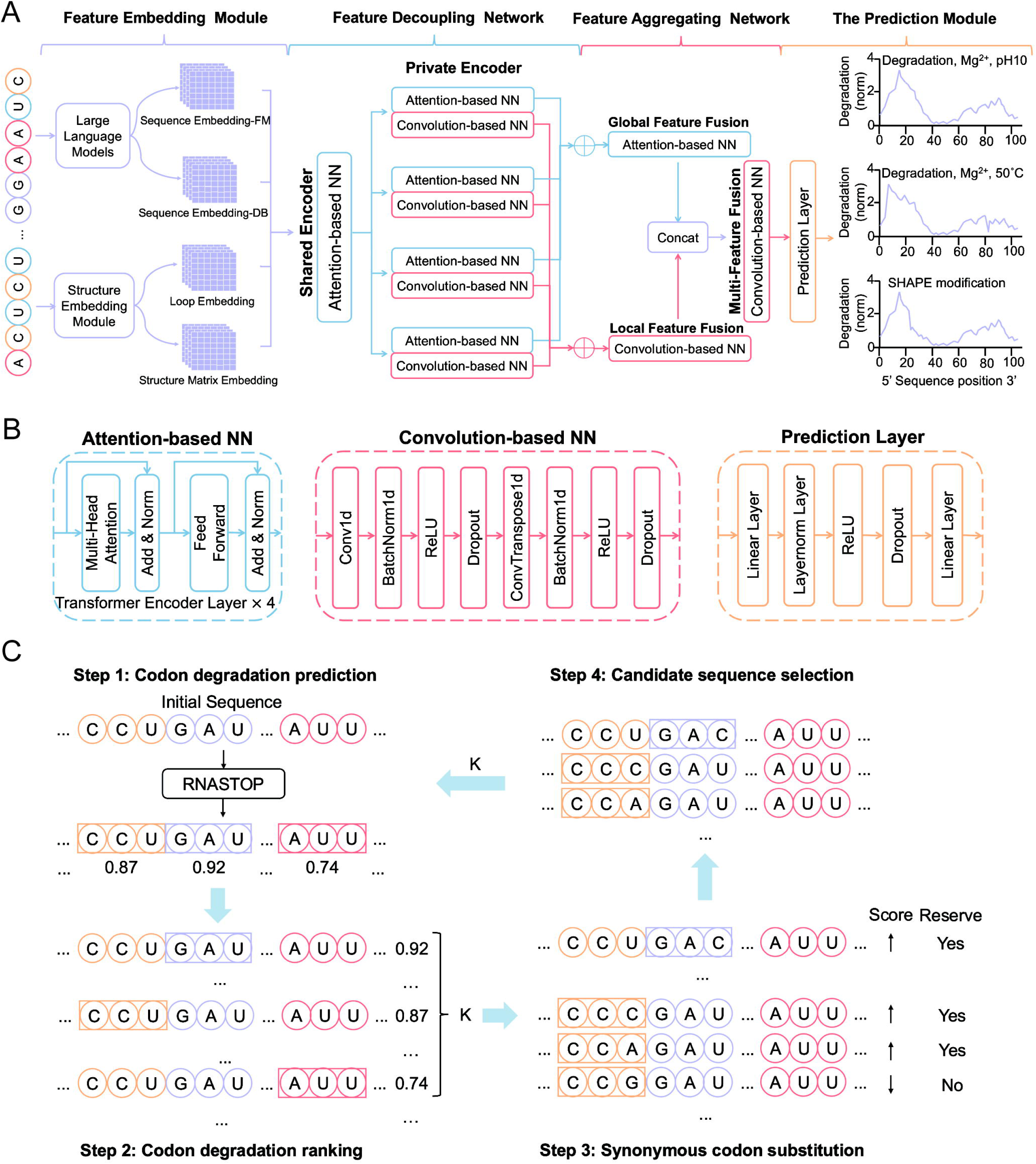
Overview of RNASTOP framework. RNASTOP encompasses an mRNA degradation prediction model and an mRNA codon optimization module. **A.** Overview of the mRNA degradation prediction model, including feature embedding module, the dual-branch feature decoupling-and-aggregating network and the prediction layer. **B.** Sub-modules of the mRNA degradation prediction model, including the attention-based neural network, the convolution-based neural network and the prediction layer. **C.** Overview of mRNA codon optimization module based on the mRNA degradation prediction model and beam search.

### Dataset

#### Stanford OpenVaccine Kaggle competition dataset

The mRNA sequences utilized in the SOVK competition were sourced from the Eterna platform [23]. The dataset comprises 3,029 mRNA sequences of length 107 nt and 3,005 mRNA sequences of length 130 nt. For the 107 nt sequences, the competition organizers provided nucleotide-level degradation profiles for the first 68 nucleotides under four accelerated degradation conditions, alongside reactivity profiles obtained via the In-line-seq method [23]: deg_pH10, deg_Mg_pH10, deg_50°C, deg_Mg_50°C, and reactivity. For the 130 nt sequences, degradation profiles for the first 102 nucleotides were provided under two conditions (deg_Mg_pH10, deg_Mg_50°C) along with reactivity profiles. To ensure data quality, these 6,034 mRNA sequences were stringently filtered based on three criteria (**Note 1 in File S1**). Following this screening process, 2,218 sequences of length 107 nt and 1,172 sequences of length 130 nt were retained. These 3,390 high-quality sequences were subsequently partitioned into a training dataset, a public test dataset, and a private test dataset (**Figure S1**).

The SOVK competition aimed to develop models capable of accurately predicting three specific target types: deg_Mg_pH10, deg_Mg_50°C, and reactivity, given the mRNA sequences and their secondary structures as inputs (**Figure 1A**) [9]. For an mRNA sequence of length *N*, the model yields 3X *N* prediction values. Model performance was evaluated using the Mean Columnwise Root Mean Squared Error (MCRMSE) across the three target types, defined as:

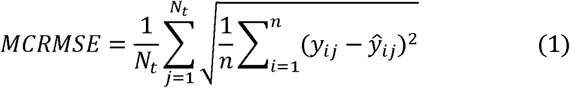

where *N_t_* is the number of scored data types, *n* is the number of nucleotides in the dataset, *y_ij_* represents the experimental ground-truth value, and *ŷ_ij_* denotes the predicted value.

##### Full-length mRNA dataset

To assess the model’s generalization capability in predicting overall degradation on full-length mRNAs, we utilized an independent dataset comprising 188 mRNAs encoding four classes of protein targets: a short multi-epitope vaccine, the model protein nano-luciferase (with one class consisting of varied untranslated regions and a second consisting of varied coding sequences), and enhanced green fluorescent protein [23]. The overall degradation rates for these full-length mRNAs were experimentally characterized using the PERSIST-seq method.

### The mRNA feature embedding module

We utilized the ViennaRNA Package [24] to compute the loop type, secondary structure, and base-pairing probability (BPP) matrix for each mRNA sequence. Secondary structures, represented in “dot-bracket” notation, were subsequently converted into adjacency and distance matrices (incorporating both first-order and second-order distances). For sequence embedding, RNASTOP incorporates representations from two nucleic acid LLMs: RNA-FM [25] and DNABERT [26] (**Notes 2, 3, and 4 in File S1**). Loop types were embedded using a standard PyTorch embedding layer. Additionally, a dedicated matrix embedding module was employed to process the BPP, adjacency, and distance matrices (**Note 5 in File S1**).

### The dual-branch feature decoupling-and-aggregating network

The dual-branch feature decoupling-and-aggregating network [27] consists of a feature decoupling module and a feature aggregating module. The feature decoupling module contains a shared encoder and four dual-branch private encoders. The shared encoder is structured as a stack of four transformer encoder blocks, encompassing multi-head self-attention mechanisms, layer normalization, and feed-forward neural networks (**Figure 1B**). The private encoder consists of a global encoder and a local encoder. The structure of the global encoders is identical to that of the shared encoder. The local encoder utilizes Conv1d, BatchNorm1d, ReLU activation, dropout, and ConvTranspose1d (**Figure 1B**). The feature aggregating module includes a global feature fusion module, a local feature fusion module, and a multi-feature fusion layer. The structure of the global feature fusion module is the same as that of the shared encoder, while the local feature fusion module and multi-feature fusion layer share the structure of the local encoder.

### Model training

The RNASTOP model was trained using the Adam optimizer [28] coupled with a cosine annealing learning rate scheduler. To maximize data utilization while enhancing the diversity of training samples, we retained 811 mRNA sequences (length 107 nt, SN_filter=0) that contained noisy labels. Early stopping was implemented to halt training if model performance on the public test set failed to improve for 50 consecutive epochs. Following hyperparameter tuning (**Figure S2, Note 6 in File S1**), we selected a learning rate of 4.0e-4, a batch size of 64, a convolutional kernel size of 7, a dropout rate of 0.3, an embedding dimension of 128, and 4 transformer encoder blocks.

### Estimation of overall full-length mRNA degradation and sequence similarity

To estimate the overall degradation of full-length mRNAs, we aggregated the single-nucleotide degradation values predicted by RNASTOP into a cumulative score:

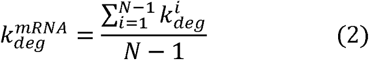

where 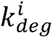 is the degradation of nucleotide linkage *i*, and *N* is the length of the mRNA. The model’s generalization performance was assessed by calculating the correlation coefficient between 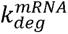 and the experimentally measured full-length mRNA degradation values. In addition, ten alternative strategies for aggregating single-nucleotide predictions were systematically evaluated (**Note 7 in File S1, Figure S3**).

Sequence similarity between mRNA constructs was quantitatively evaluated using a local sequence alignment approach, formulated as:

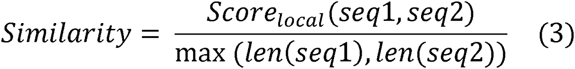

where *Score_local_* represents the optimal local alignment score computed using the *pairwise2. align. localxx* algorithm from the Biopython library. Identical nucleotides at aligned positions were awarded one point, with no penalties applied for mismatches or gaps.

### mRNA codon optimization algorithm

RNASTOP employs a beam search algorithm for mRNA codon optimization, executing four primary steps (**Figure 1C**):

Codon degradation prediction: The predictive model evaluates the degradation profile of each codon within the sequence. A codon’s overall degradation score is computed as the sum of the degradation values of its three constituent nucleotides.

Codon degradation ranking: Codons are sorted in descending order based on their degradation scores, identifying the top *K* codons most susceptible to degradation.

Synonymous codon substitution: Synonymous mutations are systematically applied to the *K* identified vulnerable codons to generate mutant sequences. Because single-codon mutations are performed iteratively, each generated sequence differs from its parent sequence by exactly one codon. Mutants demonstrating improved biophysical properties relative to the original sequence are retained as candidates.

Candidate sequence selection and iteration: Candidate sequences are evaluated and ranked based on a comprehensive scoring function:

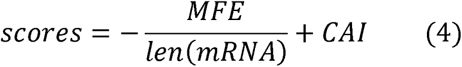

where *len(mRNA)* is the length of the mRNA sequence. The *K* sequences with the highest scores are selected for the subsequent iteration. This iterative process continues until the score converges and no further improvement is observed. The sequence attaining the highest final score is designated as the optimally designed mRNA. According to the study by Huang et al. [3], we set the beam width *K* to 500. Additionally, the performance of a strictly degradation-based scoring function was comparatively evaluated (**Note 8 in File S1, Figure S4, Table S1**).

## Results

### RNASTOP is a strong predictor of mRNA degradation

The performance of the RNASTOP model in predicting mRNA degradation at single-nucleotide resolution was evaluated on the SOVK competition dataset, which comprises both public and private test sets [9]. In this competition, 1636 teams submitted 35,806 solutions, scored using the MCRMSE. Among these submissions, the Nullrecurrent model (**Figure 2A**) ranked first, followed by the Kazuki2 model [9], where the DegScore model [23] served as the baseline (**Note 9 in File S1**). RNASTOP outperformed all 35,806 models on both the public and private test datasets, as illustrated in **Figure 2B**. On the public test set, RNASTOP achieved the lowest MCRMSE of 0.1982, outperforming both Nullrecurrent and Kazuki2 (0.2276) by approximately 0.03. More importantly, on the private test set, RNASTOP yielded an MCRMSE of 0.2962, representing a 13% improvement over the leading Nullrecurrent model (0.3420). Compared to the baseline DegScore model, the MCRMSE of RNASTOP was 49% and 37% lower on the public and private test sets, respectively. Notably, RNASTOP contains approximately 8.5 million trainable parameters, which is significantly fewer than Nullrecurrent’s 30 million, further underscoring the efficiency of the RNASTOP model. Furthermore, runtime analysis demonstrated that RNASTOP achieves an optimal balance between predictive precision and computational speed (**Note 10 in File S1, Figure S5**).

**Figure 2.**
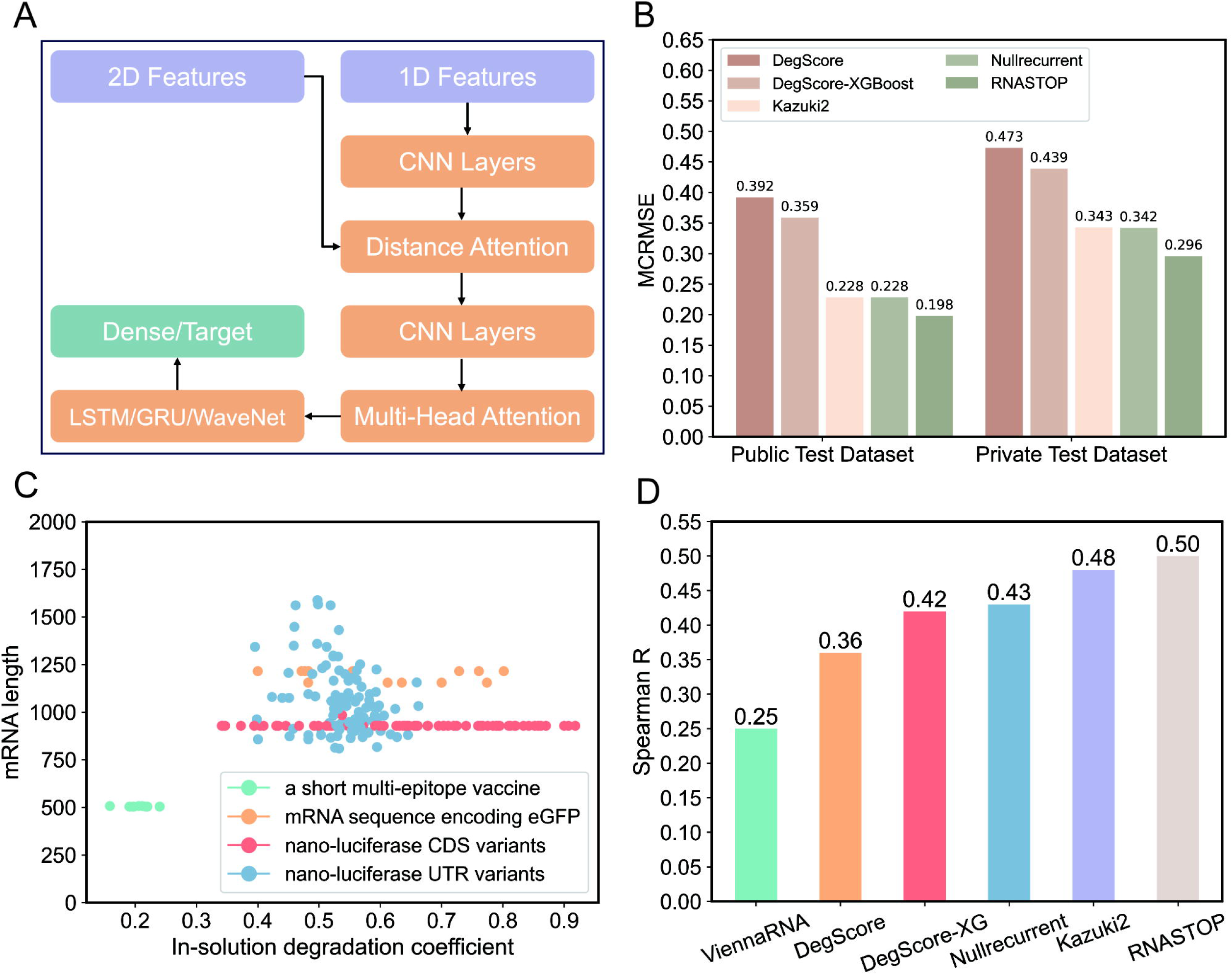
RNASTOP achieves accurate prediction of mRNA degradation. **A.** Simple schematic of the model architecture used by the Nullrecurrent. This model integrated two sets of features into the model architecture, which combined elements of classic recurrent neural networks and convolutional neural networks. **B.** RNASTOP’s MCRMSE is lower than the state-of-the-art models on both the public and private test datasets of the SOVK competition. **C.** Distribution of mRNA length and degradation coefficient on the full-length mRNA dataset. (*n*□=□188) **D.** RNASTOP’s prediction results exhibit the highest correlation to experimentally determined degradation coefficients on the full-length mRNA dataset. (number of data *n*□=□188)

Beyond single-nucleotide evaluation, we assessed RNASTOP’s capability to predict overall degradation on an independent full-length mRNA dataset [23]. The lengths of these mRNAs range from 504 to 1,588 nt (**Figure 2C**), nearly tenfold longer than the SOVK training sequences. As illustrated in **Figure 2D**, RNASTOP exhibited the highest correlation with experimentally determined overall degradation rates, achieving a Spearman correlation coefficient of 0.50 (*p*=1.7×10^-13^). The Spearman correlation coefficients for the Kazuki2 model and the Nullrecurrent model are 0.48 (*p*=3.3×10^-12^) and 0.43 (*p*=9.5×10^-10^), respectively. Notably, all three deep learning models significantly outperformed traditional unpaired probability values derived from ViennaRNA RNAfold v. 2.4.14 [24] (*R*=0.25, *p*=5.4×10^-4^), the DegScore linear regression model [23] (*R*=0.36, *p*=2.9×10^-7^), and the DegScore-XGBoost model (*R*=0.42, *p*=1.8×10^-9^). The experimental results underscore the exceptional ability of RNASTOP in accurately predicting the overall degradation of full-length mRNAs, concurrently evidencing its robust generalization capacity.

### Nucleic acid LLMs and the dual-branch architecture enable robust transfer learning

The SOVK training dataset comprises 1,189 mRNA sequences of 107 nt, whereas the private test dataset includes 629 sequences of 107 nt and 1,172 sequences of 130 nt (**Figure S1**). We visualized the sequence distributions of the training set and the private test set using *t*-distributed stochastic neighbor embedding (*t*-SNE) [29] (**Figure 3A**). The visualization revealed that the distribution of the 130-nt private test sequences significantly diverges from that of the 107-nt training sequences. Quantitative sequence alignment confirmed this, showing that the lowest sequence similarity score between the two sets was merely 0.2923. Despite this distributional shift, RNASTOP accurately predicted the degradation profiles of the 130-nt sequences at single-nucleotide resolution, demonstrating formidable transfer learning capabilities.

**Figure 3.**
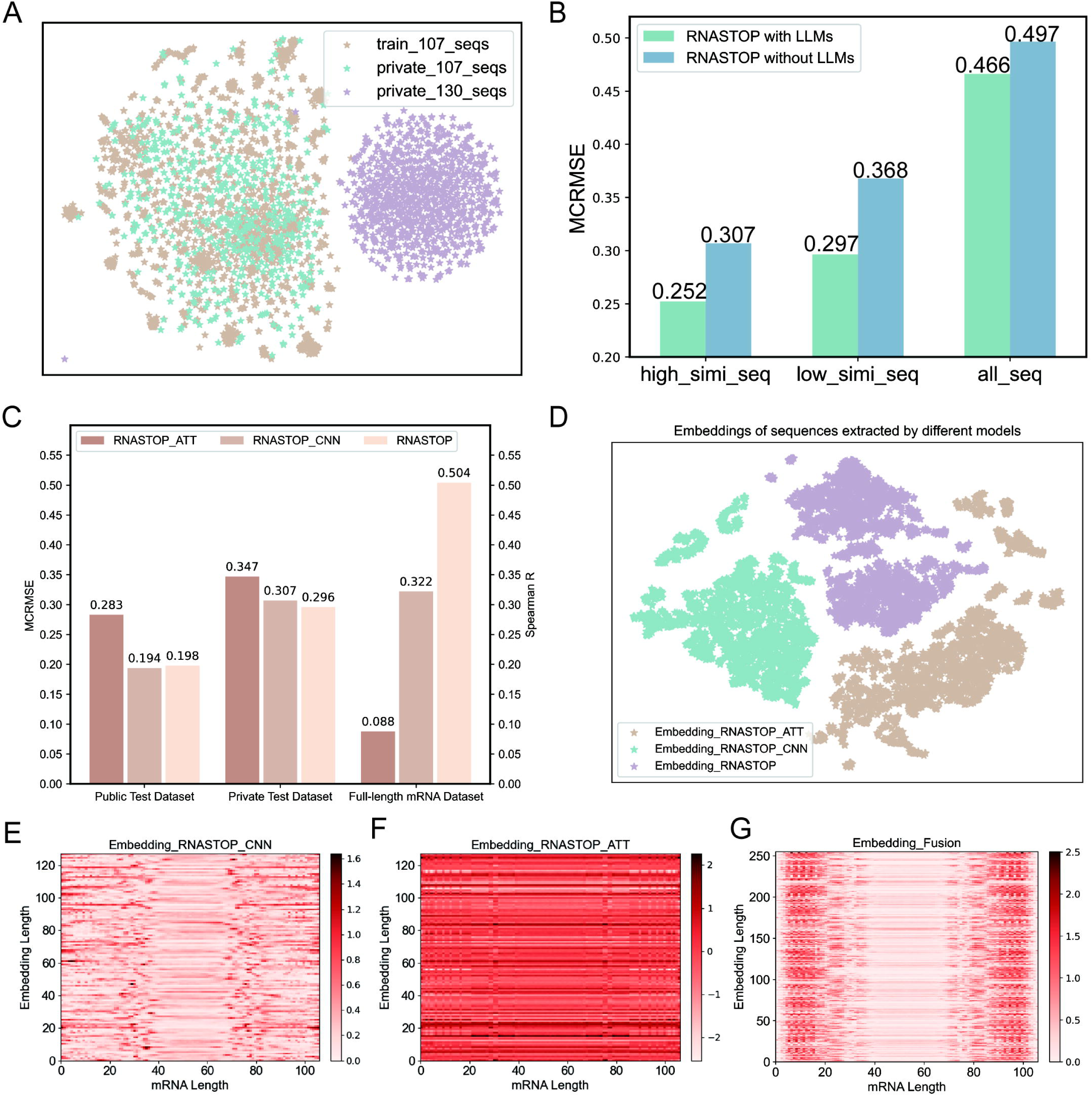
The dual-branch network architecture improves the prediction performance of RNASTOP model. **A.** mRNA distribution on the training dataset and private test dataset of the SOVK competition. (number of data points *n*□=□2,990) **B.** The nucleic acid LLMs improve the transfer learning performance of RNASTOP. **C.** The performance of different model architectures on the SOVK dataset and the full-length mRNA dataset, with MCRMSE serving as the evaluation metric for the SOVK dataset and the Spearman correlation coefficient for the full-length mRNA dataset. **D.** *t*-SNE visualization of mRNA embeddings extracted by different model architectures on the SOVK dataset. (*n*=2000) **E.** Heatmap of mRNA embeddings extracted by RNASTOP_CNN model on the SOVK dataset. (*n*=2000) **F.** Heatmap of mRNA embeddings extracted by RNASTOP_ATT model on the SOVK dataset. (*n*=2000) **G.** Heatmap of mRNA embeddings extracted by RNASTOP multi-feature fusion layer on the SOVK dataset. (*n*=2000)

To determine whether nucleic acid LLMs drive this transfer learning performance, we conducted an ablation study evaluating RNASTOP with and without nucleic acid LLM embeddings across three specific subsets of the 130-nt private test data: the sequence with the highest similarity to the training set, the sequence with the lowest similarity, and the entire set (**Figure 3B**). Across all three subsets, the nucleic acid LLM-integrated RNASTOP model demonstrated superior predictive performance. This advantage was most pronounced in the lowest-similarity subset, where the inclusion of nucleic acid LLMs reduced the MCRMSE by 0.0713 (an approximate 20% improvement). These findings confirm that pre-trained nucleic acid LLMs significantly enhance the transfer learning capabilities of the RNASTOP model.

Additionally, we investigated the contribution of the dual-branch network architecture by evaluating two ablated models: one retaining only the attention-based module (RNASTOP_ATT) and another retaining only the convolutional module (RNASTOP_CNN). As shown in **Figure 3C**, both ablated models suffered substantial performance drops on the private test dataset and the full-length mRNA dataset. On the full-length dataset, the RNASTOP_ATT model almost entirely lost its ability to predict overall degradation. This demonstrates that the dual-branch architecture is vital for the model’s transfer learning capability.

To further explore how the dual-branch network architecture improves model performance, we utilized the *t*-SNE technique to visualize the mRNA embeddings extracted by RNASTOP_ATT, RNASTOP_CNN, and RNASTOP on the SOVK dataset, as shown in **Figure 3D**. The embeddings from RNASTOP_ATT and RNASTOP_CNN occupied distinctly different latent spaces, whereas the full RNASTOP model successfully integrated both representational spaces. Heatmap visualizations of these embeddings (**Figure 3E** and **3F**) revealed that RNASTOP_CNN prioritizes specific local positional contexts (e.g., positions 0-40 and 80-100), while RNASTOP_ATT captures global attention across specific embedding dimensions (e.g., dimensions 22, 61, and 103). By concatenating these branches, the full model expands its embedding dimension from 128 to 256 (**Figure 3G**). This fused embedding effectively synthesizes information from both sub-networks, simultaneously capturing local positional motifs and critical global features. Thus, the dual-branch architecture seamlessly integrates complementary feature-extraction perspectives, equipping the RNASTOP model with robust predictive accuracy and generalization capabilities.

### RNASTOP is capable of capturing sequence and structural motifs governing degradation

To better evaluate RNASTOP’s ability to capture underlying biophysical rules, we analyzed its predictions on representative mRNA sequences. **Figures 4A-E** highlight sequences from the private test set that achieved the lowest RMSE for the deg_Mg_pH10 condition (see **Figure S6** for the top 10 sequences). The RNASTOP predictions closely mirrored the experimental ground truth, successfully identifying diverse stabilizing structural motifs. For example, paired nucleotides forming stem regions are known to exhibit high chemical stability; RNASTOP consistently predicted low degradation coefficients for these regions (**Figure 4C**). Furthermore, five-nucleotide hairpin loops are thermodynamically optimal and highly stable [30]. For the sequence in **Figure 4B**, which contains hairpins of 4, 5, and 6 nucleotides, RNASTOP accurately predicted the lowest degradation coefficient for the 5-nucleotide hairpin. Similarly, symmetrical internal loops are chemically more stable than asymmetrical ones [31], a pattern correctly identified by RNASTOP in the sequences shown in **Figures 4A** and **4E**. Conversely, **Figures 4F-G** illustrate sequences with the highest RMSE (see **Figure S7** for the top 10 worst-predicted sequences). The sequence in **Figure 4F** features an exceedingly complex secondary structure, with only 130 nucleotides yet comprising four internal loops, seven stem regions, two hairpin loops, and one bulge, posing a significant challenge for model prediction. In **Figure 4G**, the mRNA’s secondary structure includes longer single strands, with discrepancies between RNASTOP’s predictions and experimental measurements predominantly occurring in nucleotides on these single strands. The RNASTOP model tends to predict higher degradation coefficients for single-stranded nucleotides. These divergences in predicting highly complex structures (**Figure 4F**) and extended single-stranded regions (**Figure 4G**) highlight current model limitations, which are likely due to insufficient examples of these structural patterns in the training data. To address these limitations and enhance prediction accuracy, future improvements will focus on enriching the dataset with such challenging samples to better capture complex spatial dependencies.

**Figure 4.**
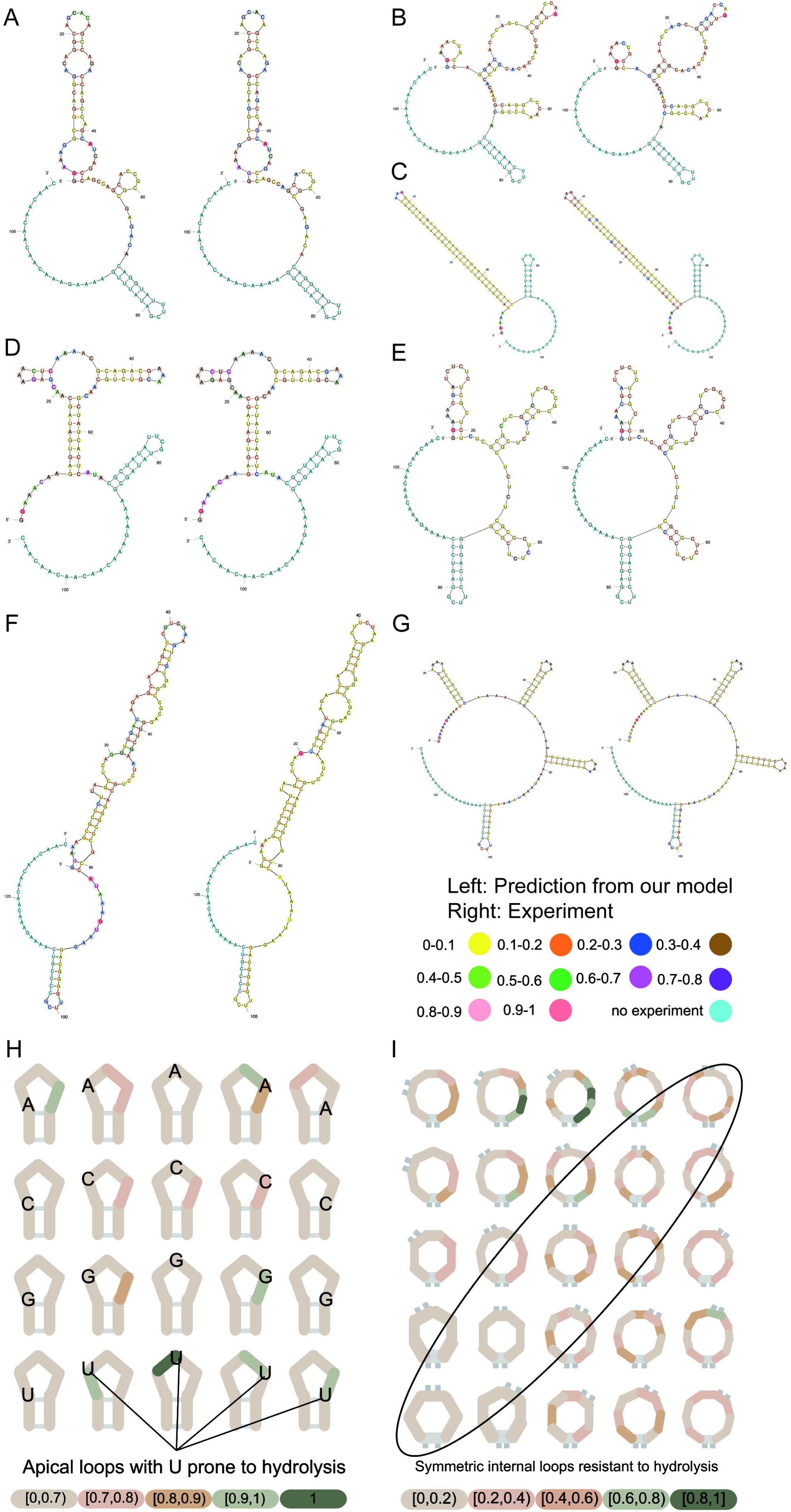
RNASTOP is capable of capturing sequence and structure patterns affecting mRNA degradation. **A-E**. Representative structures from the best-predicted mRNA sequences from degradation at 10□mM Mg2+, pH□10, 1□day, 24□°C. Different colors represent different degradation coefficients after normalization. **F-G**. Representative structures from the worst-predicted mRNA sequences from degradation at 10□mM Mg2+, pH□10, 1□day, 24□°C. **H-I.** RNASTOP reveals patterns in degradation based on both sequence (i.e., linkages ending at 3’ uridine are particularly reactive) and structure (symmetric internal loops, circled, have suppressed hydrolytic degradation compared to asymmetric internal loops). Different colors represent different degradation coefficients after normalization. The number of data are 1280 and 1000 respectively.

To further investigate whether RNASTOP captures the influence of sequence patterns on mRNA degradation, we conducted validation on 1280 mRNA sequences, with results illustrated in **Figure 4H**. Each visualization in **Figure 4H** represents the average of normalized degradation coefficients. These 1280 mRNA sequences share the same secondary structure, differing only in the types of nucleotides in their hairpin loops. Prior experimental findings have indicated that linkages that lead to a 3’ uridine are particularly prone to degradation [23]. According to the results in **Figure 4H**, RNASTOP successfully captures this trend. To delve deeper into whether RNASTOP discerns the impact of secondary structures on mRNA degradation, validation was performed on 1000 mRNA sequences, with results shown in **Figure 4I**. Each visualization in **Figure 4I** also represents the average of normalized degradation coefficients. Symmetrical internal loops are more stable than asymmetrical ones [31], and the results in **Figure 4I** suggest that the RNASTOP model also captures this trend. Furthermore, our quantitative analysis of structural motif recognition on the public test set confirmed that RNASTOP detects specific secondary structure patterns affecting mRNA degradation (**Note 11 in File S1, Figure S8**). This implies that incorporating the RNASTOP model into mRNA design algorithms would allow for an automated way to model such biochemical attributes within a designed mRNA molecule.

Additionally, we demonstrated that RNASTOP can reveal important features that affect mRNA degradation by a Leave-One-Feature-Out approach. These results suggest that mRNA sequences and BPP matrices may be key features affecting mRNA chemical stability (**Figure S9, Note 6 in File S1**).

### RNASTOP concurrently optimizes mRNA chemical stability and translation efficiency

Having established RNASTOP as an accurate and biophysically interpretable predictor, we deployed it to rationally optimize the mRNA vaccine sequences for COVID-19 and VZV. Initially, we evaluated two heuristic search algorithms: beam search and Monte Carlo tree search (MCTS). Preliminary experiments indicated that beam search was computationally more efficient (**Note 12 in File S, Figures S10 and S11**). Consequently, RNASTOP employed the beam search algorithm with a beam width of 500 for all subsequent codon optimization tasks.

Given that a lower MFE typically drives RNA into a more compact, secondary structure-rich conformation that resists in-solution degradation [23], we established MFE as a primary optimization objective. In addition to MFE, GC content, the proportion of unpaired nucleotides, and RNASTOP’s overall degradation score serve as crucial indicators of chemical stability. Furthermore, because mRNA vaccine efficacy relies on both stability and translation efficiency, we incorporated the CAI as a co-optimization objective to improve sequence translation efficiency.

Using the widely adopted BNT-162b2 COVID-19 mRNA vaccine [32] from BioNTech-Pfizer as a baseline, the progressive impact of the optimization process on RNA sequence properties is illustrated in **Figure 5A-E**. The MFE exhibited a consistent decline throughout the optimization iterations, dropping from an initial -1265.90 kcal/mol to -1531.20 kcal/mol. For comparison, state-of-the-art RNA pre-trained models, GEMORNA [18] and CodonTransformer [17], only achieved MFE reductions to -1187.6 kcal/mol and -1338.8 kcal/mol, respectively, and the commercial tool OptimumGene reached -1252.0 kcal/mol. This highlights RNASTOP’s superiority over existing methods in optimizing chemical stability. Concurrently, other chemical stability-related metrics improved: GC content increased from 0.5695 to 0.5852, the unpaired nucleotide proportion decreased from 0.3967 to 0.3443, and the overall degradation score dropped from 0.4575 to 0.4345. Crucially, the original BNT-162b2 sequence already possessed a high CAI of 0.9516, making further enhancement challenging. Nevertheless, RNASTOP still achieved a slight increase to 0.9556. In contrast, optimization by GEMORNA, CodonTransformer, and OptimumGene reduced the CAI to 0.8313, 0.8762, and 0.9049, respectively. Notably, although the LinearDesign algorithm achieved a low MFE of -2547.1 kcal/mol, its CAI decreased to 0.7325. These results confirm that RNASTOP achieves a superior and simultaneous improvement in both chemical stability and translatability.

**Figure 5.**
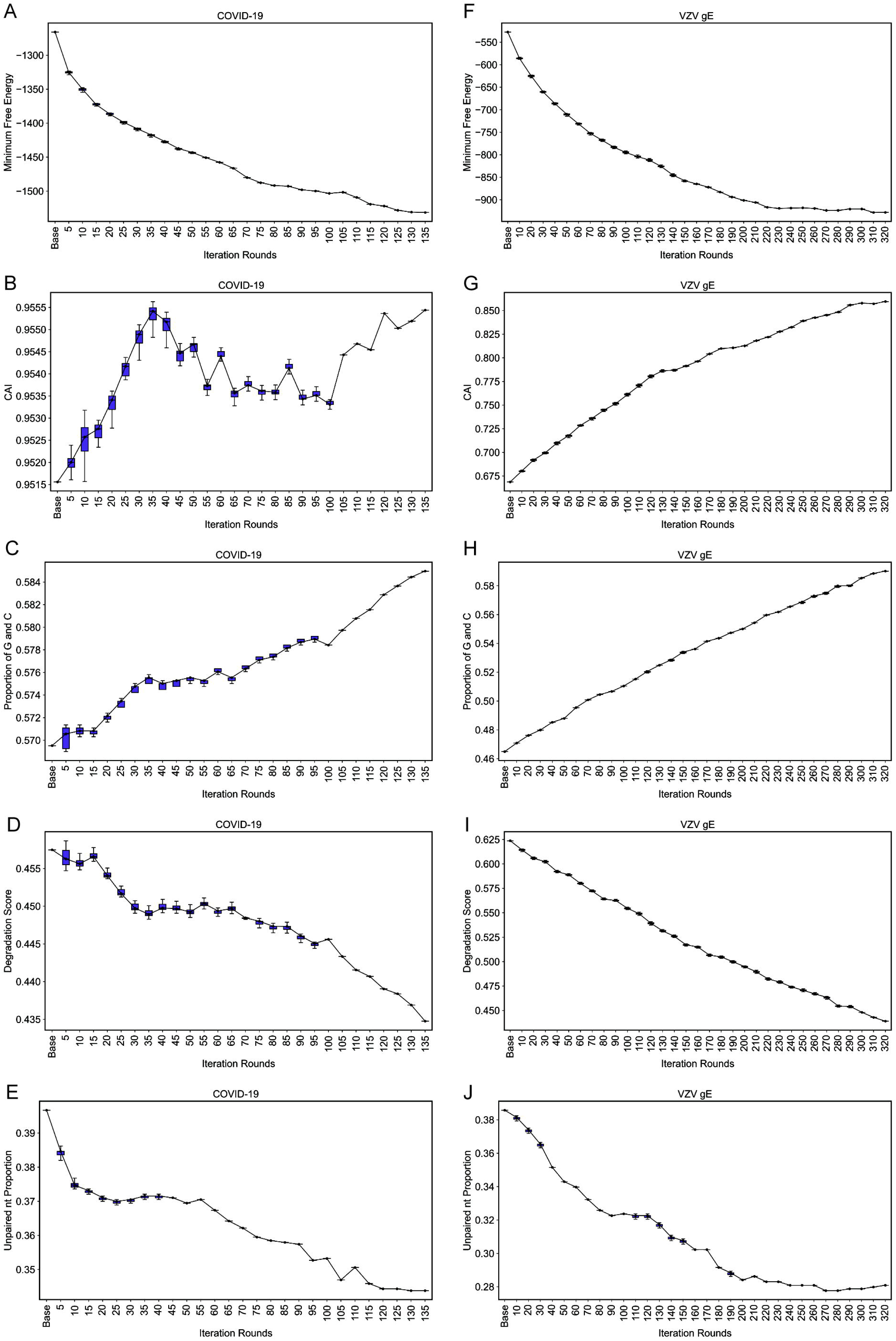
RNASTOP is capable of optimizing the mRNA sequences to enhance their chemical stability and translation efficiency. **A-E.** Changes in key metrics during the iterative optimization of the COVID-19 mRNA vaccine sequence. **F-J.** Corresponding changes for the VZV gE mRNA vaccine sequence.

To validate RNASTOP’s generalizability, we applied it to the wild-type VZV gE (glycoprotein E) mRNA vaccine sequence. As shown in **Figure 5F-J**, all evaluated metrics exhibited improvement. Following RNASTOP optimization, the MFE decreased from -527.4 kcal/mol to -926.8 kcal/mol, and the CAI increased from 0.6687 to 0.8601. Additionally, the GC content increased from 0.4650 to 0.5907, the unpaired nucleotide proportion dropped from 0.3858 to 0.2830, and the overall degradation score decreased from 0.6238 to 0.4380. Overall, RNASTOP enhanced the MFE of the COVID-19 and VZV mRNA vaccines by 20.96% and 75.73%, respectively, and improved their CAI by 0.42% and 28.62%. These findings demonstrate RNASTOP’s robust capability and broad generalizability in concurrently enhancing RNA sequence chemical stability and translation efficiency.

We further analyzed the secondary structures of the mRNA sequences before and after optimization (**Figure 6**). As shown in **Figure 6A** and **6B**, the MFE structures exhibit significant topological changes following RNASTOP optimization. The original mRNAs (left) exhibit highly branched topologies characterized by numerous single-stranded loops and short, interrupted helices. In contrast, the RNASTOP-optimized sequences (right) adopt highly compact architectures dominated by extended, continuous double-stranded stems, as highlighted in the magnified insets. This increase in base-pairing density and reduction in structural branching indicate a lower free energy state, suggesting enhanced chemical stability and potentially higher resistance to endonucleolytic degradation *in vivo*.

**Figure 6.**
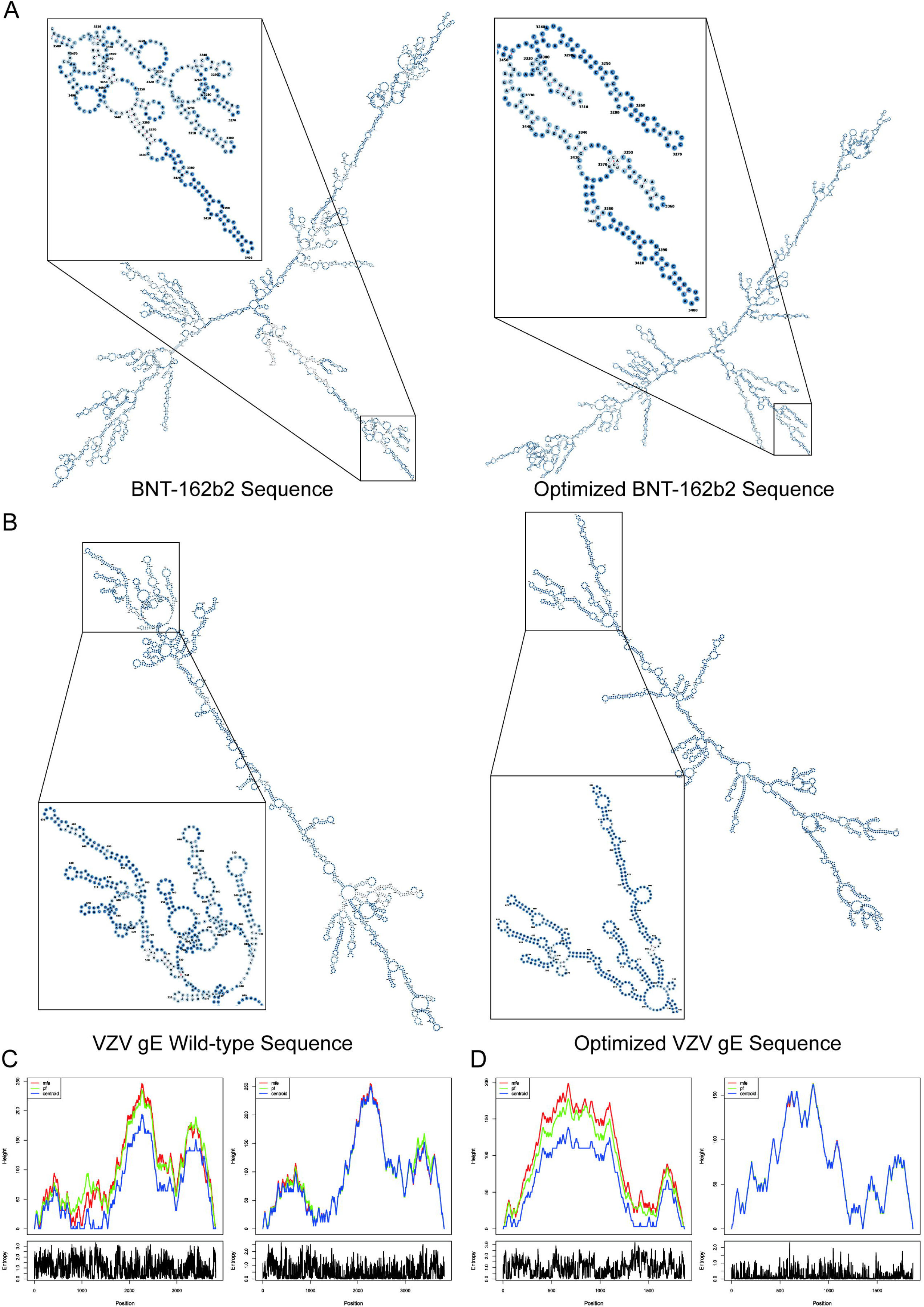
Secondary structure analysis of mRNA vaccine sequences before and after optimization. **A.** Secondary structures of BNT-162b2 (left) and the optimized sequence (right). **B.** Secondary structures of VZV gE (left) and the optimized sequence (right). **C.** Mountain plots (top) and positional entropy profiles (bottom) for BNT-162b2, comparing the original (left) and optimized (right) sequences. **D.** Corresponding plots for the VZV gE sequences, comparing the original (left) and optimized (right) versions.

This macroscopic stabilization is fundamentally driven by a reshaping of the RNA thermodynamic folding landscape, as evidenced by mountain plots and positional entropy profiles (**Figure 6C and 6D**). For the wild-type sequences, the divergence among the MFE, partition function, and centroid curves implies a complex energy landscape characterized by multiple competing sub-optimal conformations. This structural ambiguity is further reflected by the widespread high-density spikes in the positional entropy profiles, indicating significant flexibility at the single-nucleotide level. Conversely, RNASTOP optimization successfully condenses the folding trajectory into a single, well-defined global minimum. The near-perfect overlap of the MFE, partition function, and centroid curves in the optimized mountain plots, coupled with a reduction in positional entropy, demonstrates that the optimized sequences are highly deterministic. Overall, these data highlight the robust capability of RNASTOP to rationally navigate the synonymous sequence space, transforming dynamic, heterogeneous mRNA sequences into structurally uniform and biophysically stable vaccine sequences.

## Discussion

In this work, we introduce RNASTOP, an effective deep learning-based model that can accurately predict mRNA degradation and enhance mRNA chemical stability through codon optimization. The meticulously crafted network architecture empowers RNASTOP to harness the strengths of various fundamental neural network models, extracting and fusing feature embeddings of mRNAs from multiple perspectives, thereby endowing RNASTOP with superior prediction performance. By integrating nucleic acid LLMs and the dual-branch network architecture, RNASTOP is capable of predicting the degradation of mRNA sequences outside the distribution of the training set, which greatly enhances its practical utility.

Additionally, RNASTOP is capable of capturing sequence and structure patterns affecting mRNA degradation. This is a critical attribute that not only attests to its strong biological interpretability but also implies that mRNA sequences designed with RNASTOP will inherently acquire these patterns, conferring enhanced chemical stability. Moreover, we used RNASTOP to optimize mRNA vaccines for COVID-19 and VZV, demonstrating its good optimization performance and practicality.

Despite its strengths stated above, RNASTOP has some limitations. RNASTOP employs a straightforward yet effective beam search algorithm for mRNA sequence optimization. As mRNA therapeutics garner greater attention, various computational methods have been developed for mRNA sequence optimization [3, 14, 33–37], employing diverse strategies such as deep learning, lattice resolution, partition function, and quantum computing to refine mRNA sequences and enhance the effectiveness of mRNA therapeutics. In addition, RNASTOP does not account for nucleotide modifications, which impact mRNA chemical stability. With access to data on modified nucleotides for training, RNASTOP could be extended to predict and optimize the chemical stability of modified mRNA.

In conclusion, the prediction and optimization of mRNA chemical stability are pivotal to the development of mRNA therapeutics. Our research offers a novel framework, RNASTOP, that can accurately predict mRNA degradation at both single-nucleotide and full-length levels via deep learning and enhance mRNA chemical stability through codon optimization. The RNASTOP model could shorten the development time and reduce the associated costs of mRNA therapeutic development. It is a practical tool to assist in the rational design of next-generation mRNA vaccines.

## Data availability

The data used for model training and testing are also publicly available on GitHub (https://github.com/xlab-BioAI/RNASTOP) and NGDC (https://ngdc.cncb.ac.cn/biocode/tools/BT007947). This study did not generate new sequencing data.

## Code availability

The software presented in the paper is publicly available on GitHub (https://github.com/xlab-BioAI/RNASTOP). The code has also been submitted to BioCode at the National Genomics Data Center (NGDC), Beijing Institute of Genomics, Chinese Academy of Sciences / China National Center for Bioinformation (BioCode: BT007947), which is publicly accessible at https://ngdc.cncb.ac.cn/biocode/tools/BT007947.

## CRediT author statement

**Shenggeng Lin:** Investigation, Methodology, Project administration, Validation, Visualization, Writing – original draft, Writing – review & editing. **Junwei Chen:** Investigation, Writing – original draft. **Heqi Sun:** Investigation, Methodology. **Yufang Zhang:** Investigation, Methodology. **Wenyi Yang:** Visualization, Writing – review & editing. **Hong Tan:** Investigation, Methodology, Visualization. **Dong-Qing Wei:** Project administration. **Qinghua Jiang:** Project administration, Writing – review & editing. **Yi Xiong:** Investigation, Methodology, Conceptualization, Project administration, Funding acquisition, Supervision, Writing – review & editing. All authors have read and approved the final manuscript.

## Competing interests

The authors declare no competing interests.

## Acknowledgments

This work was supported by National Key Research and Development Program of China (Grant Nos. 2023YFC2506400, 2023YFC2506402 and 2024YFA1306902); National Natural Science Foundation of China (Grant No. 32570779) and Shanghai Municipal Education Commission (Grant No. 2024AIZD008). The computational experiments were partially run at the Center for High-Performance Computing, Shanghai Jiao Tong University. We thank Jianmin Wang for helpful discussions regarding the manuscript.

## Supplementary material

### File S1 Supplementary text

**Figure S1 Stanford OpenVaccine Kaggle Competition Dataset**

The composition of the Stanford OpenVaccine Kaggle competition dataset, and the division of the training dataset, the public test dataset and the private test dataset.

**Figure S2 Prediction performance of RNASTOP under different hyperparameter settings**

**Figure S3 Comparison of aggregation methods for full-length degradation prediction**

**Figure S4 Optimization effects using a scoring function based on RNASTOP’s degradation score**

**A-E.** Changes in key metrics during the iterative optimization of the COVID-19 mRNA vaccine sequence. **F-J.** Corresponding changes for the VZV gE mRNA vaccine sequence.

**Figure S5 Comparative analysis of runtime performance for RNA degradation prediction models**

**Figure S6 10 mRNA sequences with the lowest RMSE for the deg_Mg_pH10 data type out of the private test data**

**Figure S7 10 mRNA sequences with the highest RMSE for the deg_Mg_pH10 data type out of the private test data**

**Figure S8 RNASTOP captures the differential chemical stability of RNA structural motifs**

**A.** Mean predicted and experimental degradation scores by structural motif. **B.** Overall correlation between predicted and experimental degradation. **C.** Score distributions for predicted (left) and experimental (right) values.

**Figure S9 Feature importance analysis for RNASTOP mRNA degradation prediction using the leave-one-feature-out method**

**Figure S10 Optimization of mRNA vaccine sequences using beam search**

**A.** Decreasing trend of MFE values during the iterative optimization of COVID-19 and VZV mRNA vaccine sequences. In the box-and-whisker plots, the central line indicates the median, the box spans the interquartile range (IQR, 25th to 75th percentiles), and the whiskers extend to the rest of the data distribution within 1.5 × IQR. (number of data n□=□10) **B**. Specific optimization process of the COVID-19 mRNA vaccine sequence with the lowest MFE value. **C**. Specific optimization process of the VZV mRNA vaccine sequence with the lowest MFE value. **D**. Secondary structures of the initial (left) and the optimized COVID-19 mRNA vaccine sequence (right).

**Figure S11 Optimization of mRNA vaccine sequences using MCTS**

**A.** Decreasing trend of MFE values during the optimization iteration process of COVID-19 and VZV mRNA vaccine sequences. **B.** Specific optimization process of the COVID-19 mRNA vaccine sequence with the lowest MFE value. **C.** Specific optimization process of the VZV mRNA vaccine sequence with the lowest MFE value.

## Notes

### Competing Interest Statement

The authors have declared no competing interest.

## References

[1] Kramps T, Elbers K. Introduction to RNA Vaccines. Methods Mol Biol 2017;1499:1–11.

[2] Crommelin DJA, Anchordoquy TJ, Volkin DB, Jiskoot W, Mastrobattista E. Addressing the Cold Reality of mRNA Vaccine Stability. J Pharm Sci 2021;110:997–1001.

[3] Zhang H, Zhang L, Lin A, Xu C, Li Z, Liu K, et al. Algorithm for optimized mRNA design improves stability and immunogenicity. Nature 2023;621:396–403.

[4] Corbett KS, Flynn B, Foulds KE, Francica JR, Boyoglu-Barnum S, Werner AP, et al. Evaluation of the mRNA-1273 Vaccine against SARS-CoV-2 in Nonhuman Primates. N Engl J Med 2020;383:1544–55.

[5] Baden LR, El Sahly HM, Essink B, Kotloff K, Frey S, Novak R, et al. Efficacy and Safety of the mRNA-1273 SARS-CoV-2 Vaccine. N Engl J Med 2021;384:403–16.

[6] Szabo GT, Mahiny AJ, Vlatkovic I. COVID-19 mRNA vaccines: Platforms and current developments. Mol Ther 2022;30:1850–68.

[7] Zhang NN, Li XF, Deng YQ, Zhao H, Huang YJ, Yang G, et al. A Thermostable mRNA Vaccine against COVID-19. Cell 2020;182:1271–83 e16.

[8] Verbeke R, Hogan MJ, Lore K, Pardi N. Innate immune mechanisms of mRNA vaccines. Immunity 2022;55:1993–2005.

[9] Wayment-Steele HK, Kladwang W, Watkins AM, Kim DS, Tunguz B, Reade W, et al. Deep learning models for predicting RNA degradation via dual crowdsourcing. Nat Mach Intell 2022;4:1174–84.

[10] Schoenmaker L, Witzigmann D, Kulkarni JA, Verbeke R, Kersten G, Jiskoot W, et al. mRNA-lipid nanoparticle COVID-19 vaccines: Structure and stability. Int J Pharm 2021;601:120586.

[11] Kon E, Elia U, Peer D. Principles for designing an optimal mRNA lipid nanoparticle vaccine. Curr Opin Biotechnol 2022;73:329–36.

[12] Li S, Moayedpour S, Li R, Bailey M, Riahi S, Miladi M, et al. CodonBERT: Large Language Models for mRNA design and optimization. bioRxiv 2023:2023.09.09.556981.

[13] Chu Y, Yu D, Li Y, Huang K, Shen Y, Cong L, et al. A 5’ UTR Language Model for Decoding Untranslated Regions of mRNA and Function Predictions. bioRxiv 2023:2023.10.11.561938.

[14] Gong H, Wen J, Luo R, Feng Y, Guo J, Fu H, et al. Integrated mRNA sequence optimization using deep learning. Brief Bioinform 2023;24.

[15] Blakney AK. A tool for optimizing messenger RNA sequence. Nature 2023;621:262–4.

[16] Goulet DR, Yan Y, Agrawal P, Waight AB, Mak AN, Zhu Y. Codon Optimization Using a Recurrent Neural Network. J Comput Biol 2023;30:70–81.

[17] Fallahpour A, Gureghian V, Filion GJ, Lindner AB, Pandi A. CodonTransformer: a multispecies codon optimizer using context-aware neural networks. Nature Communications 2025;16:3205.

[18] Zhang H, Liu H, Xu Y, Huang H, Liu Y, Wang J, et al. Deep generative models design mRNA sequences with enhanced translational capacity and stability. Science 2025;390:eadr8470.

[19] He S, Gao B, Sabnis R, Sun Q. RNAdegformer: accurate prediction of mRNA degradation at nucleotide resolution with deep learning. Brief Bioinform 2023;24.

[20] Chen K, Zhou Y, Ding M, Wang Y, Ren Z, Yang Y. Self-supervised learning on millions of pre-mRNA sequences improves sequence-based RNA splicing prediction. bioRxiv 2023:2023.01.31.526427.

[21] Wayment-Steele HK, Kim DS, Choe CA, Nicol JJ, Wellington-Oguri R, Watkins AM, et al. Theoretical basis for stabilizing messenger RNA through secondary structure design. Nucleic Acids Res 2021;49:10604–17.

[22] Bae H, Coller J. Codon optimality-mediated mRNA degradation: Linking translational elongation to mRNA stability. Mol Cell 2022;82:1467–76.

[23] Leppek K, Byeon GW, Kladwang W, Wayment-Steele HK, Kerr CH, Xu AF, et al. Combinatorial optimization of mRNA structure, stability, and translation for RNA-based therapeutics. Nat Commun 2022;13:1536.

[24] Lorenz R, Bernhart SH, Honer Zu Siederdissen C, Tafer H, Flamm C, Stadler PF, et al. ViennaRNA Package 2.0. Algorithms Mol Biol 2011;6:26.

[25] Chen J, Hu Z, Sun S, Tan Q, Wang Y, Yu Q, et al. Interpretable RNA foundation model from unannotated data for highly accurate RNA structure and function predictions. bioRxiv 2022:2022.08. 06.503062.

[26] Ji Y, Zhou Z, Liu H, Davuluri RV. DNABERT: pre-trained Bidirectional Encoder Representations from Transformers model for DNA-language in genome. Bioinformatics 2021;37:2112–20.

[27] Zhao Z, Bai H, Zhang J, Zhang Y, Xu S, Lin Z, et al. Cddfuse: Correlation-driven dual-branch feature decomposition for multi-modality image fusion. Proceedings of the IEEE/CVF conference on computer vision and pattern recognition 2023:5906–16.

[28] Kingma DP, Ba J. Adam: A method for stochastic optimization. arXiv preprint arXiv:1412.6980 2014.

[29] Van der Maaten L, Hinton G. Visualizing data using t-SNE. Journal of machine learning research 2008;9.

[30] He Y, Shen Z, Zhang Q, Wang S, Huang D-S. A survey on deep learning in DNA/RNA motif mining. Briefings in Bioinformatics 2021;22:bbaa229.

[31] Weeks KM, Crothers DM. Major groove accessibility of RNA. Science 1993;261:1574–7.

[32] Polack FP, Thomas SJ, Kitchin N, Absalon J, Gurtman A, Lockhart S, et al. Safety and Efficacy of the BNT162b2 mRNA Covid-19 Vaccine. N Engl J Med 2020;383:2603–15.

[33] Zhang H, Sarkar A, Bertels K. A resource-efficient variational quantum algorithm for mRNA codon optimization. arXiv preprint arXiv:2404.14858 2024.

[34] Fu H, Liang Y, Zhong X, Pan Z, Huang L, Zhang H, et al. Codon optimization with deep learning to enhance protein expression. Sci Rep 2020;10:17617.

[35] Sumi S, Hamada M, Saito H. Deep generative design of RNA family sequences. Nat Methods 2024;21:435–43.

[36] Dai N, Tang WY, Zhou T, Mathews DH, Huang L. Messenger and Non-Coding RNA Design via Expected Partition Function and Continuous Optimization. arXiv preprint arXiv:2401.00037 2023.

[37] Fox DM, Branson KM, Walker RC. mRNA codon optimization with quantum computers. PLoS One 2021;16:e0259101.

