## Supplementary material for "RNASTOP: A Deep Learning Framework for mRNA Chemical Stability Prediction and Optimization": Figure_S10.pdf

A

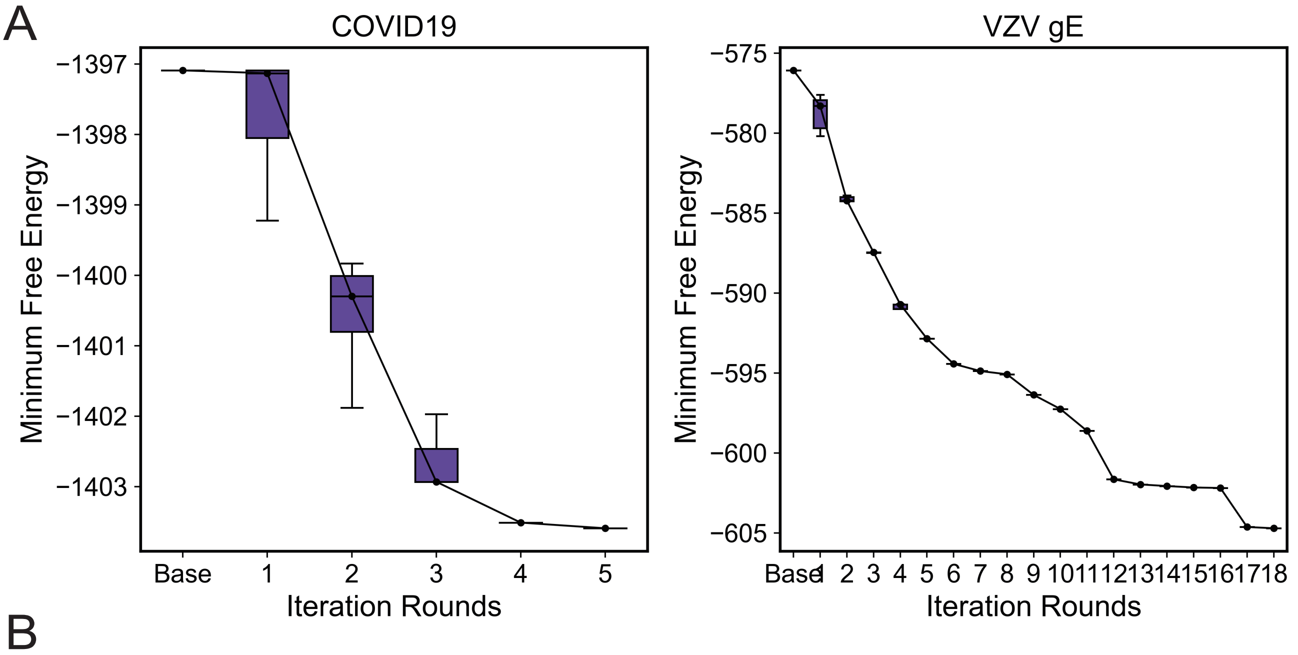


B

| Iteration round | 1 | 2 | 3 | 4 | 5 |
|---|---|---|---|---|---|
| Codon position | 642 | 855 | 1156 | 802 | 927 |
| Original codon | GUG | UUU | UUU | UUC | UUC |
| Optimized codon | GUC | UUC | UUC | UUU | UUU |

C

| Iteration round | 1 | 2 | 3 | 4 | 5 | 6 | 7 | 8 | 9 |
|---|---|---|---|---|---|---|---|---|---|
| Codon position | 221 | 147 | 557 | 551 | 92 | 88 | 555 | 276 | 581 |
| Original codon | UUA | GUA | UUA | UUA | UUA | UAU | AUA | UUA | UAU |
| Optimized codon | CUG | GUC | CUA | CUG | CUU | UAC | AUU | CUG | UAC |

| Iteration round | 10 | 11 | 12 | 13 | 14 | 15 | 16 | 17 | 18 |
|---|---|---|---|---|---|---|---|---|---|
| Codon position | 192 | 206 | 264 | 501 | 82 | 162 | 432 | 420 | 82 |
| Original codon | UAU | UUA | GUU | GUA | AUA | CUU | UGU | UUA | AUU |
| Optimized codon | UAC | CUC | GUG | GUG | AUU | UUG | UGC | CUC | AUC |

D

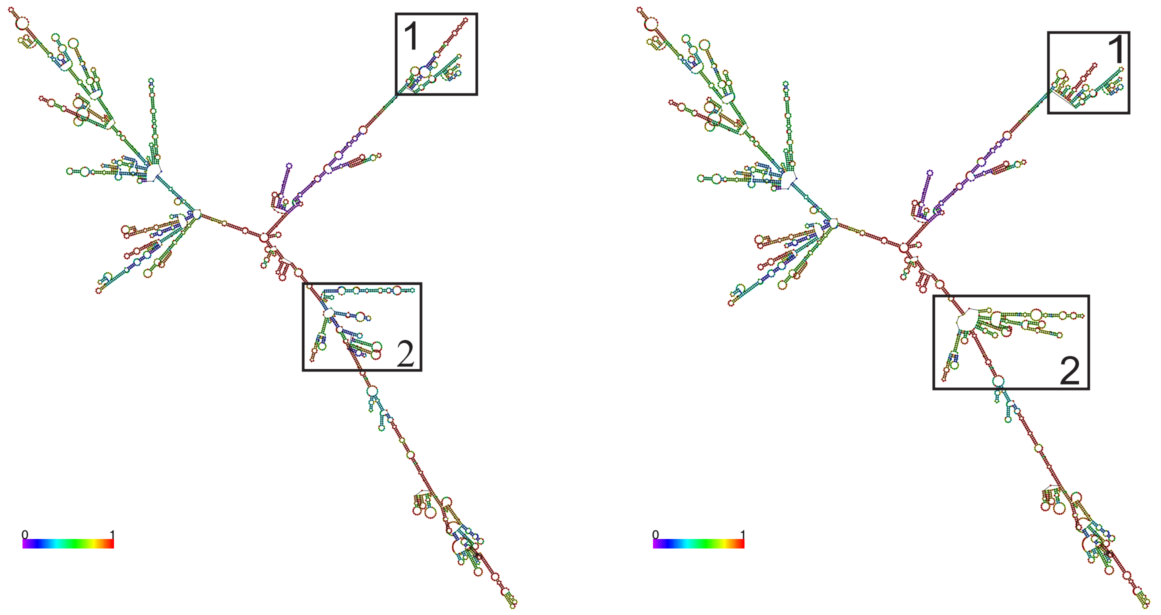
