## Supplementary material for "RNASTOP: A Deep Learning Framework for mRNA Chemical Stability Prediction and Optimization": Figure_S11.pdf

A

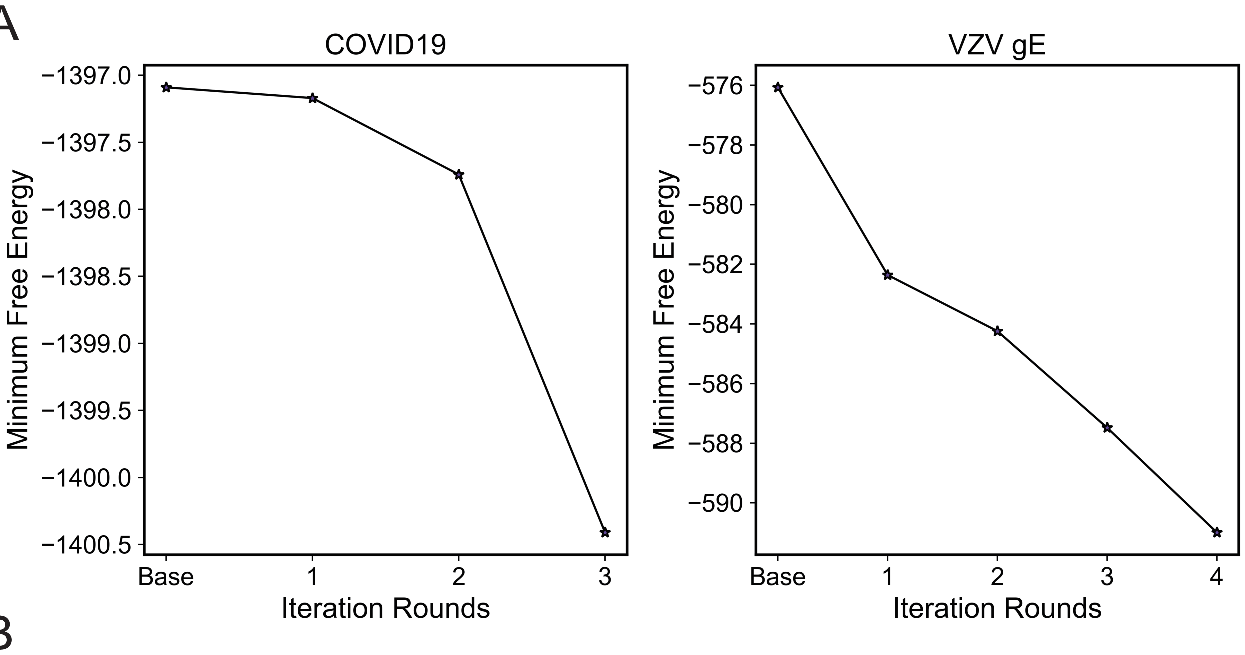


B

| Iteration round | 1 | 2 | 3 |
|---|---|---|---|
| Codon position | 927 | 802 | 642 |
| Original codon | UUC | UUC | GUG |
| Optimized codon | UUU | UUU | GUC |

C

| Iteration round | 1 | 2 | 3 | 4 |
|---|---|---|---|---|
| Codon position | 221 | 557 | 147 | 551 |
| Original codon | UUA | UUA | GUA | UUA |
| Optimized codon | CUG | CUA | GUC | CUG |
