## Supplementary material for "RNASTOP: A Deep Learning Framework for mRNA Chemical Stability Prediction and Optimization": File S1.docx

**Note 1. Stanford OpenVaccine Kaggle competition dataset**

The processing of the Stanford OpenVaccine Kaggle competition dataset is complex, hence we utilize a schematic illustration to display the data composition, filtering method, and dataset partitioning, as shown in Figure S1.

Stanford OpenVaccine Kaggle Competition Dataset screening criteria:

(i) Minimum value across all 5 profiles must be greater than -0.5;

(ii) Mean signal/noise across all 5 profiles must be greater than 1.0;

(iii) The resulting sequences were clustered into clusters with less than 50% sequence similarity, and any sequence in the test set is in a cluster that contains at most three sequences.

**Note 2. Related work on RNA large language models**

In recent years, due to the rapid development of nucleic acid large language models (LLM), several such models based on RNA sequences have also been developed [1-10]. These RNA LLMs, due to their distinct training data, varying pre-training tasks, and fine-tuning methods, exhibit respective strengths in different downstream tasks. Specifically, certain RNA LLMs, including RNA-FM [1], RNABERT [2], RNA-MSM [3], Uni-RNA [4] and CodonBERT [5], have shown remarkable capabilities in the prediction of RNA structures and functionalities. RNA-FM [1], pioneering among RNA LLMs, significantly improved the accuracy of RNA structure and functionality predictions through pre-training on 23 million RNA sequences. RNABERT [2] and RNA-MSM [3] have broadened the scope of applications to include complex downstream tasks, such as RNA family clustering and one-dimensional solvent accessibilities tasks. Uni-RNA [4] has achieved state-of-the-art performance on multiple RNA structure and function prediction tasks by pre-training on the largest RNA sequence dataset to date. Distinctively, CodonBERT [5] adopts codons as the basic unit of input, achieving codon-level representations via pre-training on over 10 million RNA sequences. RNAErnie [10] elevates pretraining by integrating RNA motifs as biological priors and introduces a type-guided fine-tuning strategy, exhibiting its superiority in both supervised and unsupervised learning.

In parallel, other models like SpliceBERT [6] and BigRNA [7] have been developed with a focus on RNA splicing modeling. SpliceBERT [6] advances the modeling of sequence-based RNA splicing by generating high-dimensional representations that encapsulate nucleotide evolutionary information alongside functional features of splicing sites. Celaj et al. developed BigRNA [7] to predict splicing by training on thousands of genome-matched datasets. Furthermore, the untranslated regions (UTR) are also crucial for RNA expression, prompting some RNA LLMs to focus on these regions, such as UTR-LM [8] for 5’UTRs and 3UTRBERT [9] for 3’UTRs. The UTR-LM performed excellently in predicting mean ribosome loading, translation efficiency, and mRNA expression level. And the 3UTRBERT demonstrated remarkable predictive capabilities in predicting RNA-protein binding sites, m6A RNA modification sites, and RNA sub-cellular localizations.

**Note 3. The RNA-FM large language model**

RNA-FM is an RNA LLM proposed by Li et al. The RNA-FM framework comprises a stacked architecture of 12 transformer encoder blocks, originally proposed in BERT [11]. Each encoder block consists of a feed-forward layer with a hidden size of 640 and a 20-head self-attention layer. Layer normalization [12] and residual connections [13, 14] are applied before and after every block, respectively. During the pre-training phase, the authors employed a self-supervised training paradigm akin to that of BERT. Approximately 15% of nucleotide tokens were randomly replaced with a special mask token. Specifically, if the i-th token was selected, the authors replaced it with the [MASK] token 80% of the time, with a random token 10% of the time, and retained the original i-th token 10% of the time. The authors then trained the model with masked language modeling by predicting the original masked tokens to minimize the cross-entropy loss. This training strategy can be formalized as the following objective function:

$$\mathcal{L}_{MLM}=\mathbb{E}_{x\sim X}\mathbb{E}_{x_{\mathcal{M}}\sim x}\sum_{i\in\mathcal{M}} -\log p(x_{i}|x_{\mathcal{/M}}) (1)$$

where a set of indices $\mathcal{M}$ is randomly sampled from each of the input sequences $x$ (comprising 15% of the entire sequence). The sequence is then masked by replacing the true token at each index $i$ with a mask token. Next, given the masked sequence ($x_{\mathcal{/M}}$) as context, the objective function minimizes the negative log-likelihood of the corresponding true nucleotide $x_{i}$. Minimizing this objective function enables the model to capture dependencies between the masked tokens and the unmasked segments within the input sequence, thereby facilitating precise predictions for the masked positions. Consequently, training RNA-FM via this formulation enables the network to learn rich, contextual representations of each sequential token.

The large-scale dataset used for the pre-training phase of RNA-FM is sourced from RNA-central [15], the most extensive non-coding RNA dataset to date [16]. This dataset aggregates non-coding RNA sequences sourced from 47 different databases, amounting to approximately 27 million RNA sequences. To curtail redundancy while preserving the dataset’s breadth, the authors employed cd-hit-est [17] to eliminate identical sequences. This process resulted in a comprehensive dataset comprising around 23.7 million ncRNA sequences. Detailed information regarding RNA-FM can be referenced in the literature [1].

**Note 4. The DNABERT large language model**

DNABERT is a DNA-focused LLM proposed by Davuluri and colleagues [18]. DNABERT follows the same training process as BERT. DNABERT first takes a set of sequences represented as k-mer tokens as input. Each sequence is represented as a matrix $M$ by embedding each token into a numerical vector. Formally, DNABERT captures contextual information by performing the multi-head self-attention mechanism on $M$:

$$MultiHead\left( M \right)=ConCat\left( {head}_{1},\ldots,{head}_{h} \right)W^{o} (2)$$

where

$${head}_{i}=softmax\left( \frac{MW_{i}^{Q}MW_{i}^{K^{T}}}{\sqrt{d_{k}}} \right)\times MW_{i}^{V} (3)$$

$W^{O}$ and $W_{i}^{Q}$, $W_{i}^{k}$, $W_{i}^{V}$ are learned parameters for linear projection. $head$ calculates the next hidden states of $M$ by first computing the attention scores between every two tokens and then utilizing them as weights to sum up lines in $MW_{i}^{V}$. $MultiHead()$ concatenates results of $h$ independent $head$ with different set of $[W_{i}^{Q}$, $W_{i}^{k}$, $W_{i}^{V}]$. The entire procedure is performed $L$ times with $L$ being number of layers.

Similar to BERT, DNABERT adopts a pre-training and fine-tuning scheme. The authors, however, made modifications to the pretraining process originally implemented in BERT by omitting next sentence prediction, adjusting the sequence length, and compelling the model to predict contiguous k tokens tailored to the DNA context. During pre-training, DNABERT learns basic syntax and semantics of DNA through self-supervision, leveraging sequences ranging from 10 to 510 in length extracted from the human genome via truncation and sampling methods. For each sequence, the authors randomly mask regions of k contiguous tokens that constitute 15% of the sequence and let DNABERT to predict the masked sequences based on the remainder, ensuring ample training examples. The authors pre-trained DNABERT with cross-entropy loss:

$$L=\sum_{i=0}^{N} -y_{i}^{'}log(y_{i}) (4)$$

here, $y_{i}^{'}$ and $y_{i}$ are the ground-truth and predicted probability for each of $N$ classes. The pre-trained DNABERT model can be fine-tuned with task-specific training data for applications in various sequence- and token-level prediction tasks. Detailed information regarding DNABERT can be referenced in the literature [18].

**Note 5. The matrix embedding module**

We employ a matrix embedding module to embed the base-pairing probability (BPP) matrices, the adjacency matrices, and distance matrices. The shapes of both the BPP matrix and the adjacency matrix are [sequence length, sequence length, 1], whereas the first-order and second-order distance matrices have shapes of [sequence length, sequence length, 3]. Initially, we concatenate the BPP matrix, the adjacency matrix, and distance matrices into a structural matrix of shape [sequence length, sequence length, 8]. This structural matrix is then embedded into a matrix of dimensions [sequence length, sequence length, 128] using the matrix embedding module. The matrix embedding module comprises a Conv2d layer, BatchNorm2d layer, ReLU layer, Dropout layer, and AdaptiveAvgPool2d layer. Detailed descriptions of these layers can be found on the PyTorch official website (https://pytorch.org/docs/stable/index.html).

**Note 6. RNASTOP reveals important features in RNA degradation prediction**

The hyperparameters of the RNASTOP model may impact its performance. To set reasonable values for the model’s hyperparameters, we discuss the influence of six hyperparameters on model performance: learning rate, batch size, size of the convolutional kernel, dropout rate, embedding dimension, and the number of layers of the transformer encoder block. According to the experimental results, the learning rate has a significant impact on model performance. If the learning rate is greater than 10^-4^, the performance of the model will decrease significantly. Other hyperparameters have a less pronounced effect on performance. Ultimately, we selected a learning rate of 4.0e-4, a batch size of 64, a convolutional kernel size of 7, a dropout rate of 0.3, an embedding dimension of 128, and a transformer encoder block layer number of 4.

To ascertain the important features for predicting mRNA degradation, we employed a straightforward Leave-One-Feature-Out approach. We retrained our models from scratch with the best settings but left out one of the available features (sequences, BPP matrixes, predicted loop types, distance matrixes, or nucleic acid LLMs), and then assessed model performance in the absence of each feature [19]. The exclusion of sequence data resulted in the most pronounced deterioration in model performance (Figure S9). This finding suggests that the mRNA sequence may be an important feature affecting mRNA degradation. Therefore, using more stable sequence motifs or motif pairs in mRNA design may improve the chemical stability of mRNA [20]. This also highlights the capability of RNASTOP in sequence motif recognition. The performance after excluding sequence information is followed by BPPs, which depict the array of potential folds an mRNA sequence can assume. This experimental result highlights the role of BPPs as an accurate representation of mRNA secondary structure[21]. In addition, this experimental result also proves that RNASTOP learns 2D feature patterns such as BPPs. The impact of the nucleic acid LLMs on performance is primarily evident on the private test set, underscoring the enhancement in the model’s generalization capability conferred by the nucleic acid LLMs. Omitting the distance matrixes and loop features has little impact on performance. The minimal impact of distance matrixes and loop features is consistent with prior findings on the significance of BPPs [21]. These features represent only the single most likely fold based on BPPs and therefore do not to portray mRNA secondary structure as comprehensively as BPPs do. These experimental results demonstrate that RNASTOP can reveal important features for mRNA degradation prediction.

**Note 7. Comparison of aggregation methods for full-length degradation prediction**

Ten different methods for combining single-nucleotide predictions to estimate the degradation of a full-length mRNA were tested. These methods included the mean (sum of scores divided by sequence length), sum, median, maximum, minimum, standard deviation, variance, 25th percentile, 75th percentile, and 90th percentile. For each method, the Spearman rank correlation coefficient between the combined prediction score and the actual experimental degradation value was calculated.

The results show that the mean method performed the best (Figure S3), achieving a Spearman correlation of 0.5039 (p < 1.69×10⁻¹³). The 75th percentile method ranked second with a correlation of 0.4937 (p < 6.05×10⁻¹³). The max method performed the worst and was not statistically significant (0.1307, p = 0.0738). These findings indicate that the mean method is the best way to convert nucleotide-level predictions into a single full-length score. However, it should be noted that this simple averaging method is still an approximation, because it does not account for the complex, non-additive interactions between local structures and neighboring nucleotides. To achieve more accurate predictions, training a model directly on a large dataset of full-length degradation scores would likely be a better approach.

**Note 8. Evaluation of a degradation-based scoring function for sequence optimization**

During beam search optimization, the default minimum free energy (MFE)-based scoring function for each RNA sequence is -MFE/len(mRNA) + CAI. In addition, we explored the optimization effect of a scoring function based on RNASTOP’s degradation score, which is -deg_sum/len(mRNA) + CAI, where deg_sum is the total predicted degradation score of the sequence. The optimization results are shown in Figure S4.

For both the COVID-19 and Varicella-Zoster Virus (VZV) mRNA vaccines, the degradation-based scoring function improved all evaluated metrics. As the optimization rounds progressed, CAI, GC content, and degradation scores steadily improved. However, as detailed in Table S1, the MFE-based scoring function performed better across all metrics. For the COVID-19 mRNA vaccine, the MFE-based function improved the MFE, CAI, and degradation score by 20.96%, 0.42%, and 5.03%, respectively. In contrast, the degradation-based function only improved them by 1.71%, 0.23%, and 3.10%. Similar trends were observed for the VZV mRNA vaccine, where the MFE-based function improved the same metrics by 75.73%, 28.62%, and 29.79%, while the degradation-based function improved them by 11.74%, 13.06%, and 13.87%. Thus, the MFE-based scoring function clearly produces better optimization results.

The main strength of RNASTOP is its ability to provide nucleotide-resolution degradation scores, thus guiding the mutation positions in heuristic search. However, selecting the best final sequence requires a reliable overall score. RNASTOP estimates the overall score simply by averaging the individual nucleotide scores. In contrast, MFE is a widely accepted and rigorous physical metric for measuring the overall stability of a mRNA sequence. Consequently, using MFE in the scoring function yields superior optimization outcomes.

**Note 9. Detailed description of the Nullrecurrent, Kazuki2, and DegScore models**

*Nullrecurrent*

The architecture of the winning Nullrecurrent model depicts the architecture of this shared kernel, which feeds 1D and 2D features, including adjacency matrices based on the secondary structure of the RNA inputs, into a multi-head attention network, the output of which is then fed into convolutional neural net layers. Additionally, the participants use sophisticated methods to process training data:

1. The participants did not use the SN_filter. Instead, they assigned NaN values to individual positions with large errors (started with error > 10 and value/error < 1.5 - and diversity can be added by varying this). The participator edited the loss function so that the Nan targets won’t contribute to loss during training. About 20K values are labeled as Nan across all 5 targets.

2. They calculated the edit distance between sequences, and did clustering based on it (trying to reverse engineer what the organizer did to the data). They found that many clusters only contain 1 sequence, but some clusters contain as many as 60 sequences. Thus, they decided to give sample weights proportional to 1/sqrt(count_in_cluster).

3. The participants assigned column weights [0.3,0.3,0.3,0.05,0.05] to favor the scored columns

Additionally, Nullrecurrent model uses data augmentation and pseudo-labeling. More detailed information about Nullrecurrent model can be found https://www.kaggle.com/c/stanford-covid-vaccine/discussion/189620.

*Kazuki2*

The architecture of Kazuki2 demonstrates an example implementation of using pseudo-labelling. The practice of pseudo-labelling, which is similar to the student-teacher learning paradigm, involves using predictions from one model as “mock ground truth” labels for another model.

In fact, the architecture of Kazuki2 is more like a stack of some deep learning models, including the gate recurrent unit neural network, the long short-term memory neural network, the graph neural network and XGBoost. More detailed information about Kazuki2 model can be found https://www.kaggle.com/competitions/stanford-covid-vaccine/discussion/189709.

*DegScore*

DegScore models degradation at a given nucleotide i as a linear function of nucleotides surrounding i:

$$Y_{i}=\sum_{k=-w}^{w} [\sum_{n\in\{A,C,G,U\}} (\beta_{k,n}I_{i+k,n})]+\sum_{k=-w}^{w} [\sum_{s\in\{H,E,I,M,B,S\}} (\beta_{k,s}I_{i+k,s})]+\beta_{0} (5)$$

where $\beta$ represents learned coefficients and $I$ is an indicator function corresponding to the identity of nucleotide $i+k$. $I$ accounts for sequence identity n (A, C, G, U) and its secondary structure type assignment s (S = stem, E = external loop, I = internal loop, B = bulge, H = hairpin, M = multiloop). $w$ is the maximum window distance. The DegScore linear regression model is available at [www.github.com/eternagame/DegScore](http://www.github.com/eternagame/DegScore).

**Note 10. Runtime and Computational Efficiency Analysis**

To evaluate computational efficiency, the runtime of RNASTOP was compared with the two fastest tools: DegScore and DegScore-XG. All tests were conducted on a single machine under identical hardware and software configurations (CPU: Intel Xeon Gold 6530, 15 cores; GPU: NVIDIA GeForce RTX 4090; RAM: 100 GB; consistent CUDA environment). Total inference times were measured across two datasets: the public test set (400 sequences) and the private test set (1,801 sequences).

As a simple linear regression model, DegScore expectedly exhibited the shortest inference times on both the public (0.27 s) and private (1.58 s) datasets (Figure S5). Between the two more complex models, RNASTOP processed the public set faster than DegScore-XG (0.93 s vs. 1.09 s), while taking slightly longer on the larger private set (6.47 s vs. 5.66 s). The slightly longer runtime of RNASTOP is an acceptable trade-off considering its higher prediction accuracy. Furthermore, processing 400 sequences in under one second and 1,801 sequences in under seven seconds demonstrates that RNASTOP is highly capable of supporting large-scale RNA analyses. Overall, the model achieves an optimal balance between predictive precision and computational efficiency.

**Note 11. Quantitative analysis of structural motif recognition**

To evaluate the ability of RNASTOP to recognize structural motifs that affect mRNA chemical stability, the entire public test set (400 mRNA sequences) was analyzed. The RNASTOP model was used to predict degradation scores for every nucleotide position. Based on secondary structure, each nucleotide was labeled by its motif type: Stem, External loop, Hairpin loop, Internal loop, Multi-loop, Bulge or Other. Predicted degradation scores were grouped by motif type and compared with experimental values.

The results show that RNASTOP accurately identifies how different structural motifs affect mRNA chemical stability (Figure S8A). Seven motif types were analyzed, with sample sizes ranging from 568 to 14,492 nucleotides. For all motif groups, predicted values correlated strongly with experimental values (Pearson r = 0.697 to 0.901, all p < 0.001; R² = 0.466 to 0.811, Figure S8B). The External loop motif showed the highest correlation (r = 0.901, R² = 0.811), followed by the Hairpin loop motif (r = 0.853, R² = 0.723). Furthermore, statistical tests confirmed that the model can distinguish the chemical stability of different structures. The difference in predicted degradation scores was especially clear between the Stem motif and all other motifs (Figure S8C). For example, base-paired Stem regions had much lower mean degradation scores (0.331) than unpaired regions like External loops (0.705) and Hairpin loops (0.588), which matches the known fact that stem regions are more stable. When comparing the mean degradation scores across all 21 possible motif pairs, 18 pairs showed statistically significant differences (using a standard threshold of p < 0.05), and 15 of these pairs showed highly significant differences (p < 0.001). These data confirm that RNASTOP successfully recognizes and measures the impact of different structural motifs on mRNA chemical stability.

**Note 12. Preliminary experimental results of codon optimization for improving mRNA chemical stability based on beam search and Monte Carlo tree search algorithms**

In order to compare the efficiency of beam search and Monte Carlo tree search (MCTS) algorithms, we conducted preliminary experiments. In the preliminary experiment, we chose the beam size of beam search to be 10 and ensured that the running times of the MCTS and beam search algorithms were equal.

The beam search-based optimization algorithm has been described in the main text. The MCTS algorithm comprises four distinct steps: selection, expansion, simulation, and backpropagation. Within this codon optimization framework, each node in the tree corresponds to a specific mRNA sequence.

Selection: MCTS calculates the Upper Confidence Bound (UCB) value for each child node, opting to expand the one possessing the highest UCB value. The UCB value is calculated as follows：

$${UCB}_{i}=\frac{Q_{i}}{N_{i}}+C\sqrt{\frac{lnN_{p}}{N_{i}}} (6)$$

where $Q_{i}$ signifies the total value of the $i$ child node，$N_{i}$ represents the number of times the $i$ child node has been visited, $N_{p}$ represents the number of times the parent node has been visited, and $C$ is a constant, typically selected to be 2.

Expension: Select the $K$ codons most susceptible to degradation for synonymous mutation through the mRNA degradation prediction model. Then select the K mutation sequences with the lowest MFE value among the obtained mutation sequences and incorporate these as child nodes within the tree.

Simulation: Calculate the MFE value corresponding to the child node and utilize the inverse as the simulation value for the child node.

Backpropagation: Upon completion of the simulation, MCTS backpropagates the simulation value of the child node backward to the root node. This process involves updating both the total revenue and the visitation count for each node as follows:

$$Q_{i}\leftarrow Q_{i}+v (7)$$

$$N_{i}\leftarrow N_{i}+1 (8)$$

where $v$ represents the simulation value of the child node.

The probability of the MCTS algorithm finding the optimal policy increases with the number of iterations. To facilitate a comparative analysis between the performance of the MCTS algorithm and that of the beam search technique in the preliminary experiment, we ensured that the execution times for both the MCTS and beam search algorithms were equivalent. Additionally, the number of child nodes expanded by MCTS was consistent with the beam width.

The specific process of optimizing COVID-19 and VZV mRNA vaccine sequences using beam search is depicted in Figure S10. The optimization process’s impact on the MFE value of the COVID-19 mRNA vaccine sequence is illustrated in Figure S10A, with specific optimization steps depicted in Figure S10B. Notably, the MFE value exhibited a consistent decline throughout the optimization process, particularly when UUU codons at positions 855 and 1156 were substituted with UUC codons, leading to a significant MFE reduction. This alteration is attributed to changes in the vaccine sequence’s secondary structure, as evidenced in Figure S10D. After five optimization iterations, the MFE value of the vaccine sequence diminished by 6.5 kcal/mol.

The decreasing trend of the MFE values of the VZV mRNA vaccine sequence and the specific optimization process are depicted in the Figure S10A and Figure S10C, respectively. The MFE value of the sequence has significantly decreased in the 1st, 2nd, 3rd, 4th, 12th, and 17th optimization iterations. A common characteristic of these iterations was the diminution of adenine (A) and uracil (U) nucleotides, coupled with an increase in cytosine (C) and guanine (G) nucleotides within the sequence. Following eighteen optimization iterations, the MFE value of the VZV vaccine sequence was reduced by 28.63 kcal/mol.

MCTS is a search algorithm based on random simulation, which increases the probability of finding the optimal strategy as the number of simulations increases. Theoretically, with an ample volume of simulations, MCTS is capable of identifying the optimal solution. However, considering the efficiency of sequence optimization algorithms is also crucial in the vaccine design process. Therefore, to limit the running time of MCTS and to facilitate comparative analysis of the performance between MCTS and beam search algorithms, we ensured that the execution times for both MCTS and beam search algorithms were equal. The specific process of optimizing COVID-19 and VZV mRNA vaccine sequences using MCTS is depicted in Figure S11. Within the same operational timeframe, the MCTS algorithm performed 3 and 4 iterative optimizations for the COVID-19 and VZV mRNA vaccine sequences, respectively. For the COVID-19 mRNA vaccine sequence, after three rounds of optimization, MCTS and beam search algorithms reduced its MFE values by 3.32 kcal/mol and 5.84 kcal/mol, respectively. For the VZV mRNA vaccine sequence, after four rounds of optimization, both MCTS and beam search algorithms achieved a reduction of 14.92 kcal/mol in its MFE value. Consequently, this also demonstrates that beam search can find more stable sequences than MCTS when optimizing for the same number of rounds.

**References:**

[1] Chen J, Hu Z, Sun S, Tan Q, Wang Y, Yu Q, et al. Interpretable RNA foundation model from unannotated data for highly accurate RNA structure and function predictions. bioRxiv 2022:2022.08. 06.503062.

[2] Akiyama M, Sakakibara Y. Informative RNA base embedding for RNA structural alignment and clustering by deep representation learning. NAR Genom Bioinform 2022;4:lqac012.

[3] Zhang Y, Lang M, Jiang J, Gao Z, Xu F, Litfin T, et al. Multiple sequence alignment-based RNA language model and its application to structural inference. Nucleic Acids Res 2024;52:e3.

[4] Wang X, Gu R, Chen Z, Li Y, Ji X, Ke G, et al. UNI-RNA: UNIVERSAL PRE-TRAINED MODELS REVOLUTIONIZE RNA RESEARCH. bioRxiv 2023:2023.07.11.548588.

[5] Li S, Moayedpour S, Li R, Bailey M, Riahi S, Miladi M, et al. CodonBERT: Large Language Models for mRNA design and optimization. bioRxiv 2023:2023.09.09.556981.

[6] Chen K, Zhou Y, Ding M, Wang Y, Ren Z, Yang Y. Self-supervised learning on millions of pre-mRNA sequences improves sequence-based RNA splicing prediction. bioRxiv 2023:2023.01.31.526427.

[7] Celaj A, Gao AJ, Lau TTY, Holgersen EM, Lo A, Lodaya V, et al. An RNA foundation model enables discovery of disease mechanisms and candidate therapeutics. bioRxiv 2023:2023.09.20.558508.

[8] Chu Y, Yu D, Li Y, Huang K, Shen Y, Cong L, et al. A 5’ UTR Language Model for Decoding Untranslated Regions of mRNA and Function Predictions. bioRxiv 2023:2023.10.11.561938.

[9] Yang Y, Li G, Pang K, Cao W, Li X, Zhang Z. Deciphering 3’ UTR mediated gene regulation using interpretable deep representation learning. bioRxiv 2023:2023.09.08.556883.

[10] Wang N, Bian J, Li Y, Li X, Mumtaz S, Kong L, et al. Multi-purpose RNA language modelling with motif-aware pretraining and type-guided fine-tuning. Nature Machine Intelligence 2024:1-10.

[11] Devlin J, Chang M-W, Lee K, Toutanova K. Bert: Pre-training of deep bidirectional transformers for language understanding. arXiv preprint arXiv:1810.04805 2018.

[12] Ba JL, Kiros JR, Hinton GE. Layer normalization. arXiv preprint arXiv:1607.06450 2016.

[13] He K, Zhang X, Ren S, Sun J. Identity mappings in deep residual networks. Computer Vision–ECCV 2016: 14th European Conference, Amsterdam, The Netherlands, October 11–14, 2016, Proceedings, Part IV 14 2016:630-45.

[14] He K, Zhang X, Ren S, Sun J. Deep residual learning for image recognition. Proceedings of the IEEE conference on computer vision and pattern recognition 2016:770-8.

[15] RNAcentral 2021: secondary structure integration, improved sequence search and new member databases. Nucleic acids research 2021;49:D212-D20.

[16] Amin N, McGrath A, Chen Y-PP. Evaluation of deep learning in non-coding RNA classification. Nature Machine Intelligence 2019;1:246-56.

[17] Fu L, Niu B, Zhu Z, Wu S, Li W. CD-HIT: accelerated for clustering the next-generation sequencing data. Bioinformatics 2012;28:3150-2.

[18] Ji Y, Zhou Z, Liu H, Davuluri RV. DNABERT: pre-trained Bidirectional Encoder Representations from Transformers model for DNA-language in genome. Bioinformatics 2021;37:2112-20.

[19] He S, Gao B, Sabnis R, Sun Q. RNAdegformer: accurate prediction of mRNA degradation at nucleotide resolution with deep learning. Brief Bioinform 2023;24.

[20] Leppek K, Byeon GW, Kladwang W, Wayment-Steele HK, Kerr CH, Xu AF, et al. Combinatorial optimization of mRNA structure, stability, and translation for RNA-based therapeutics. Nat Commun 2022;13:1536.

[21] Wayment-Steele HK, Kim DS, Choe CA, Nicol JJ, Wellington-Oguri R, Watkins AM, et al. Theoretical basis for stabilizing messenger RNA through secondary structure design. Nucleic Acids Res 2021;49:10604-17.
