## Supplementary material for "RNASTOP: A Deep Learning Framework for mRNA Chemical Stability Prediction and Optimization": Table S1.docx

**Table S1 Comparison of optimization effects using different scoring functions**

|  | **COVID-19 mRNA vaccine** | | | **VZV mRNA vaccine** | | |
| --- | --- | --- | --- | --- | --- | --- |
|  | Initial sequence | MFE-based scoring function | Degradation score-based scoring function | Initial sequence | MFE-based scoring function | Degradation score-based scoring function |
| MFE (kcal/mol) | -1265.90 | -1531.20 | -1287.60 | -527.4 | -926.8 | -589.3 |
| CAI | 0.9516 | 0.9556 | 0.9538 | 0.6687 | 0.8601 | 0.7560 |
| GC content | 0.5695 | 0.5852 | 0.5750 | 0.4650 | 0.5907 | 0.5056 |
| Degradation score | 0.4575 | 0.4345 | 0.4433 | 0.6238 | 0.4380 | 0.5373 |
| Unpaired nucleotide proportion | 0.3967 | 0.3443 | 0.3873 | 0.3858 | 0.2830 | 0.3804 |
