## Supplementary figures and images for "RNASTOP: A Deep Learning Framework for mRNA Chemical Stability Prediction and Optimization"

### Figure_S1.pdf

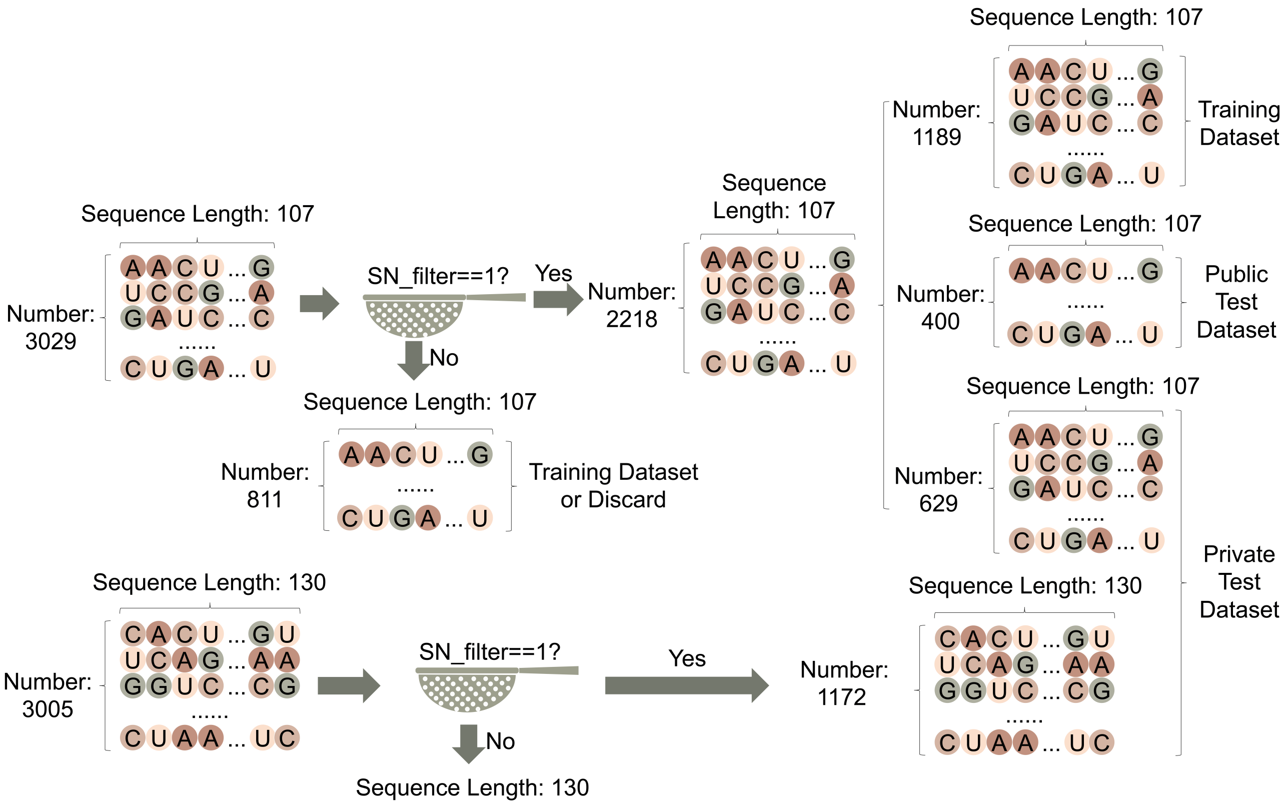

### Figure_S2.pdf

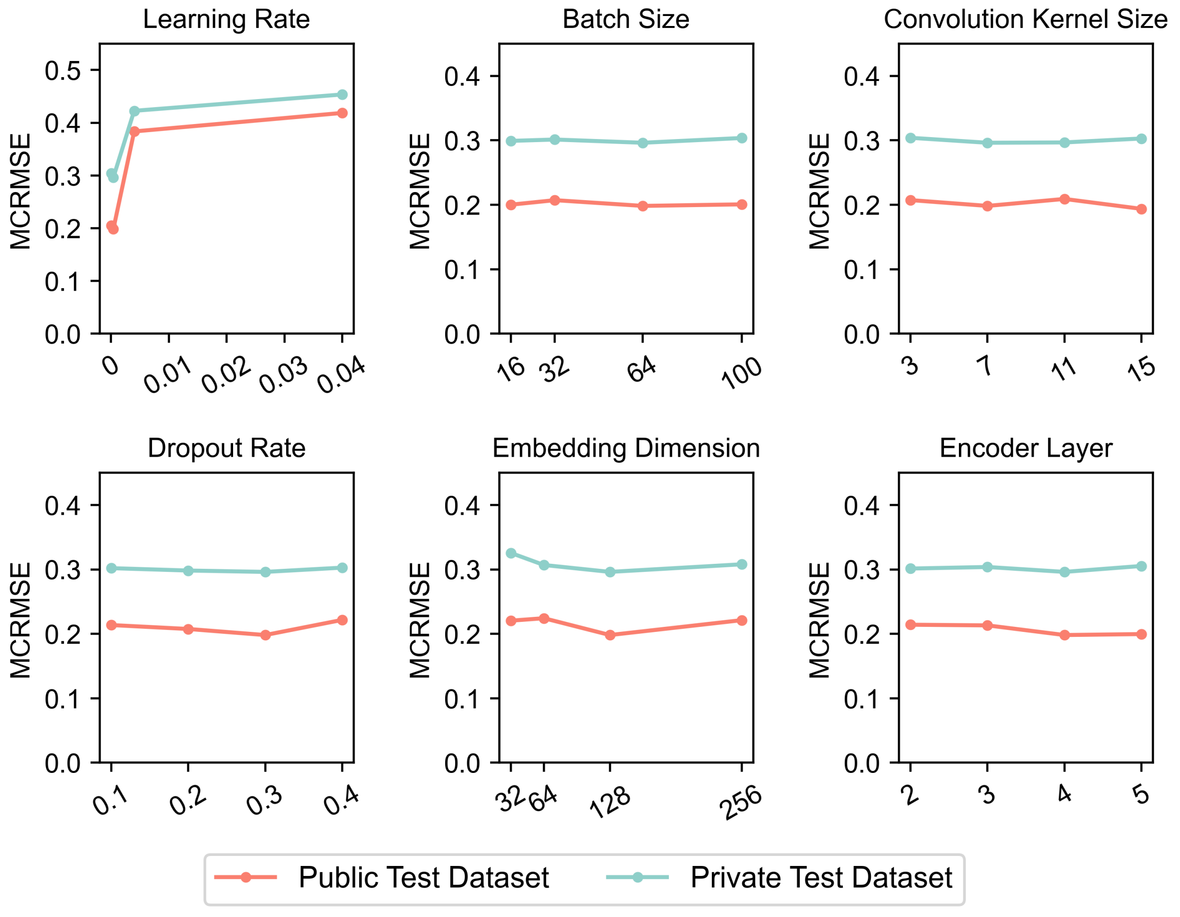

### Figure_S3.pdf

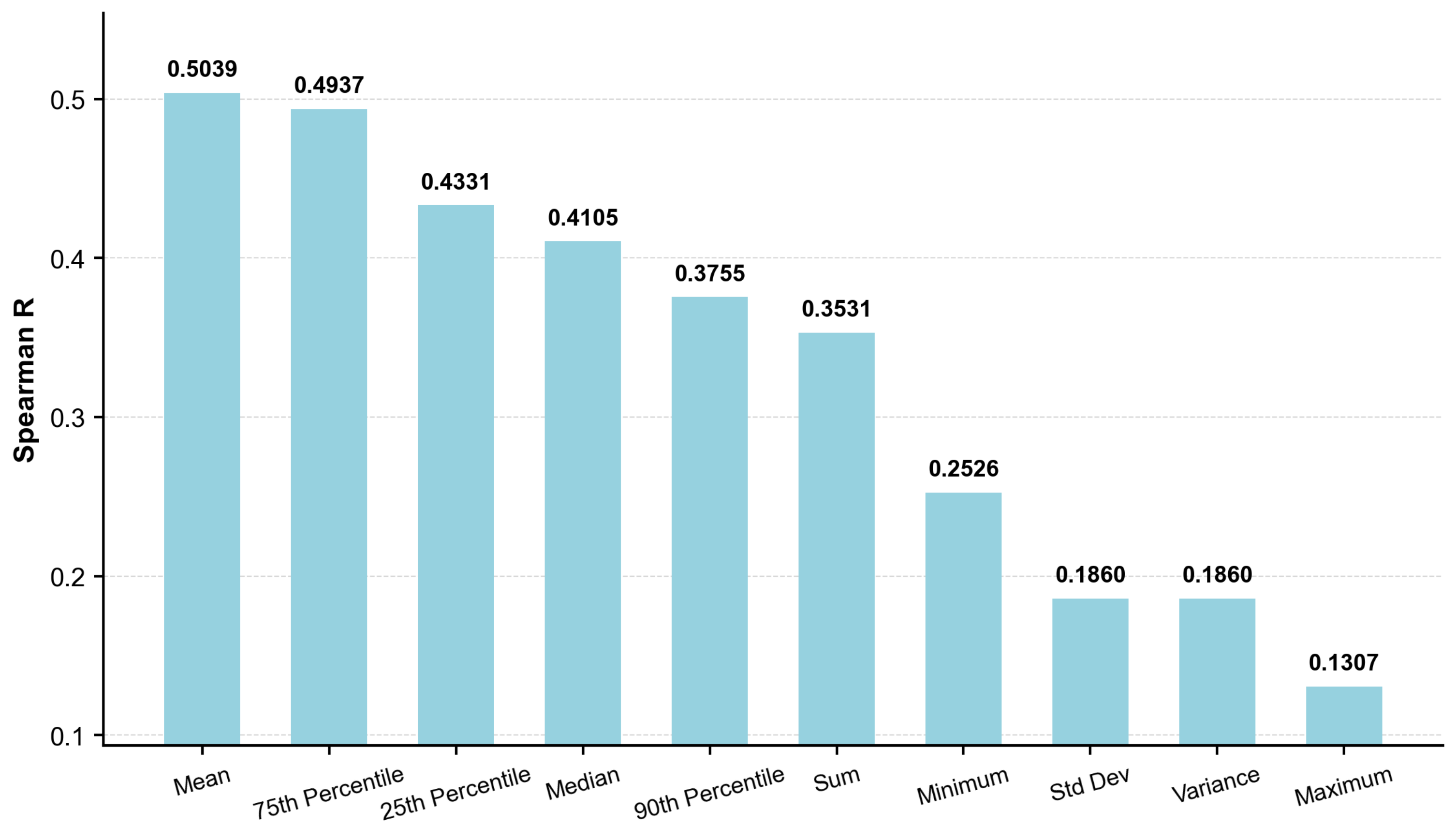

### Figure_S4.pdf

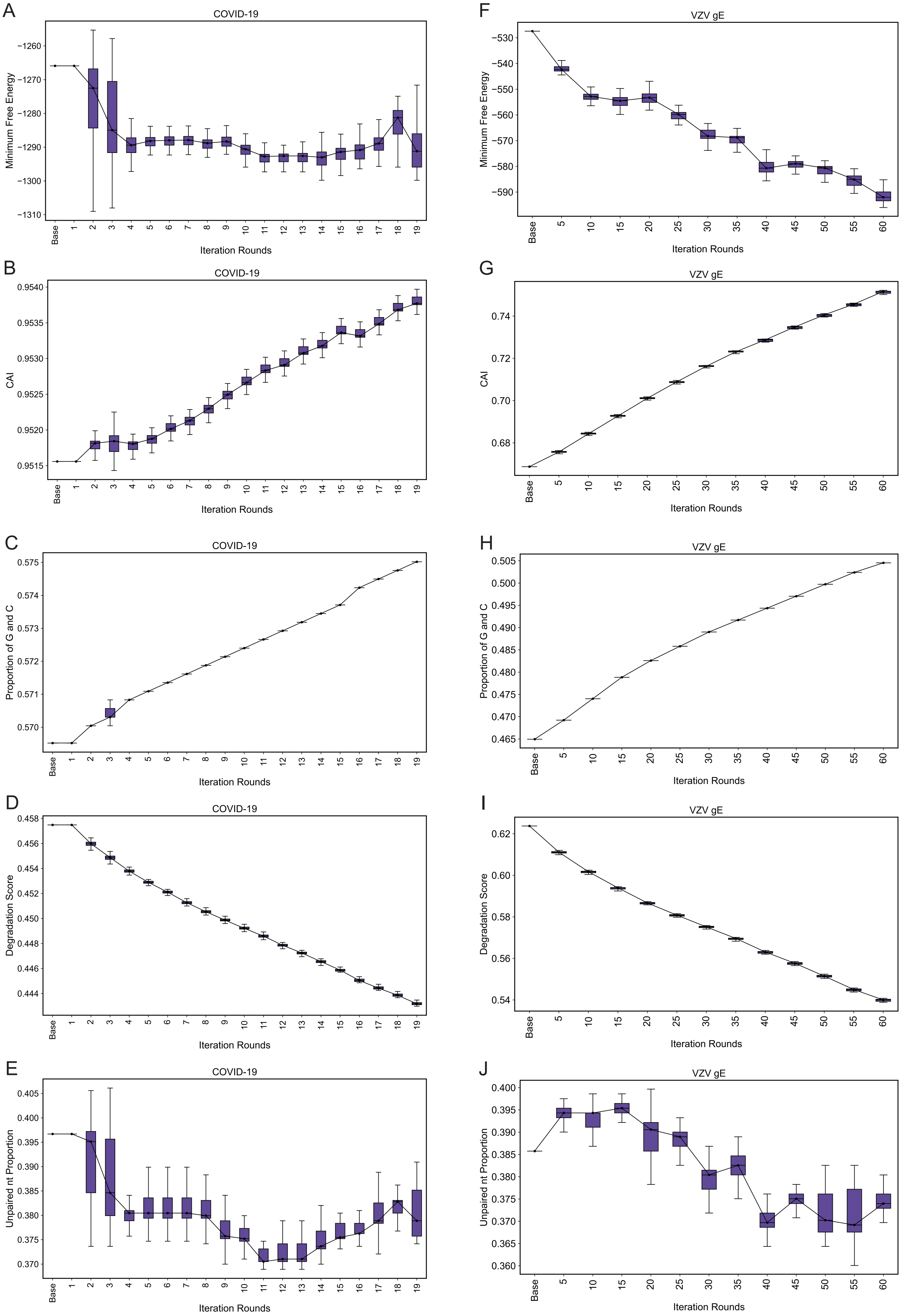

### Figure_S5.pdf

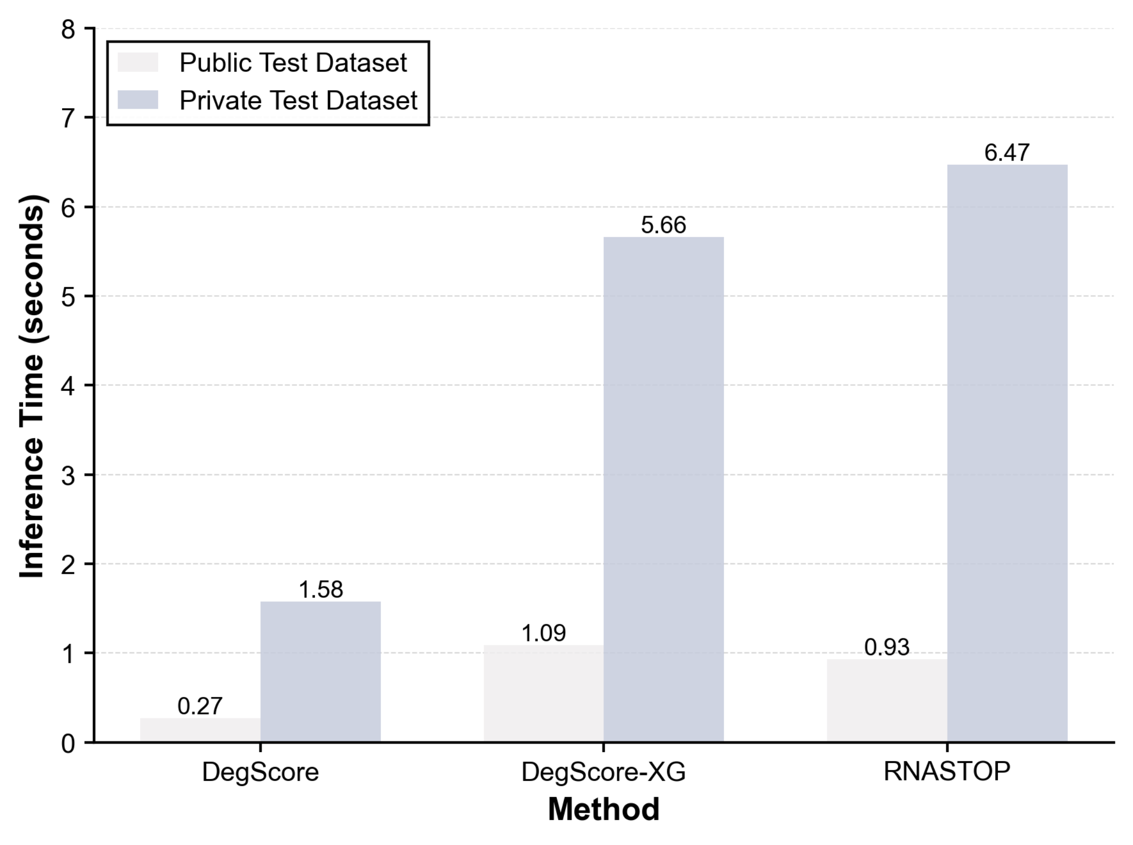

### Figure_S6.pdf

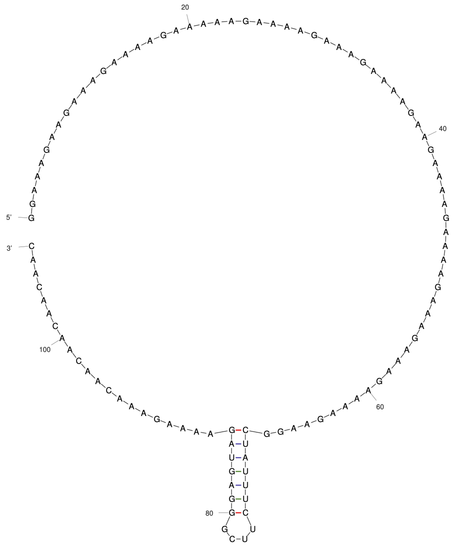

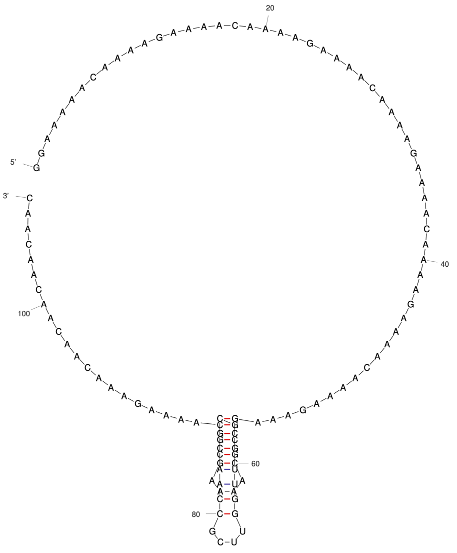

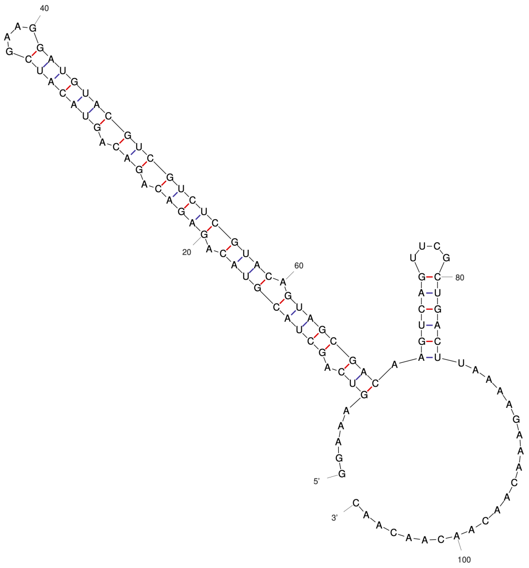

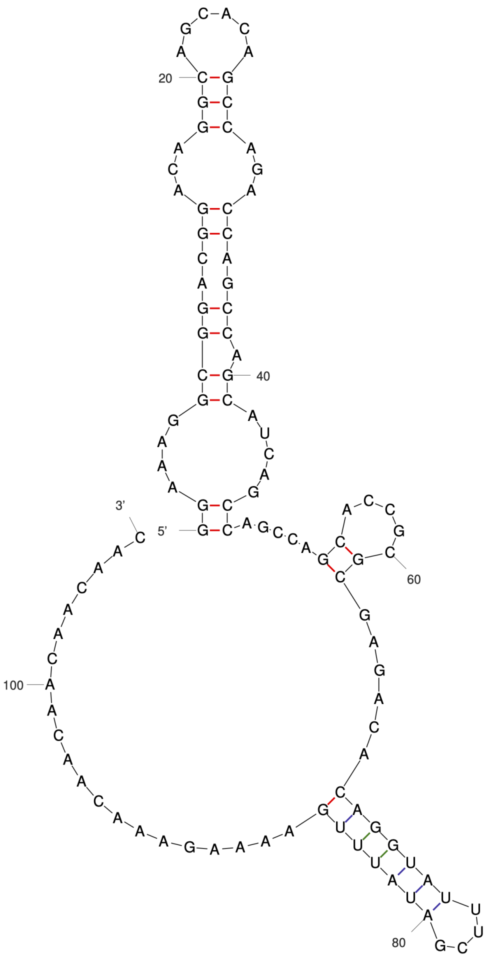

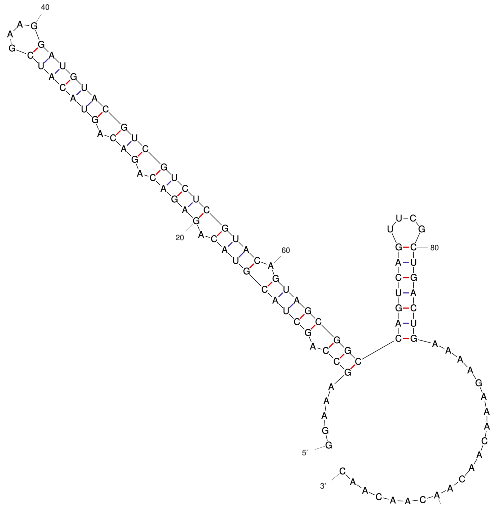

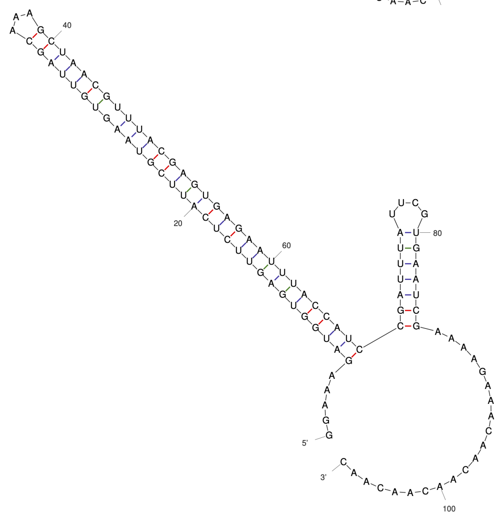

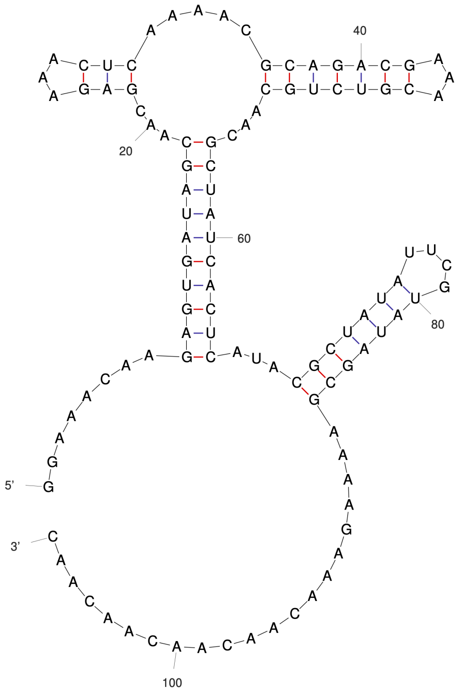

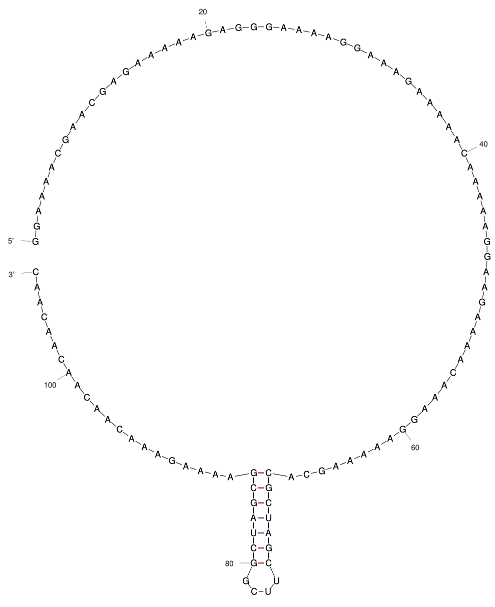

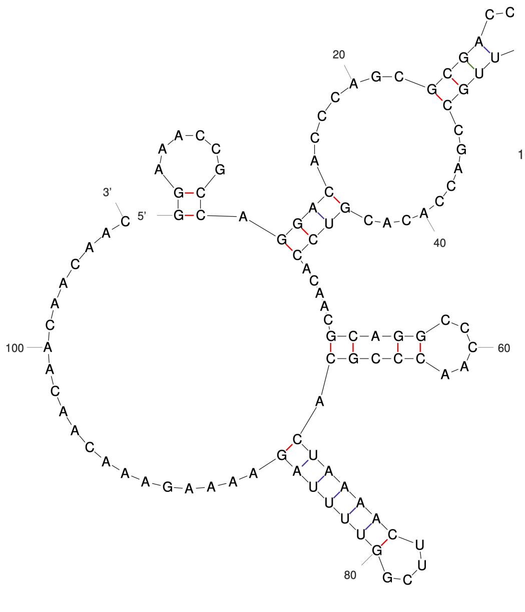

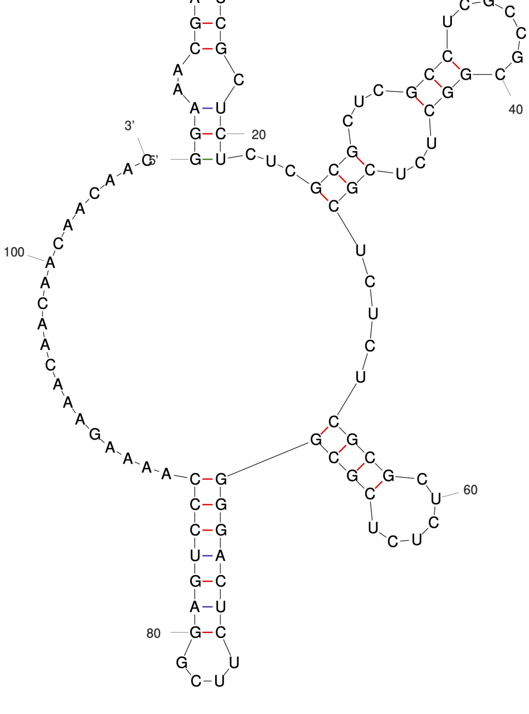

### Figure_S7.pdf

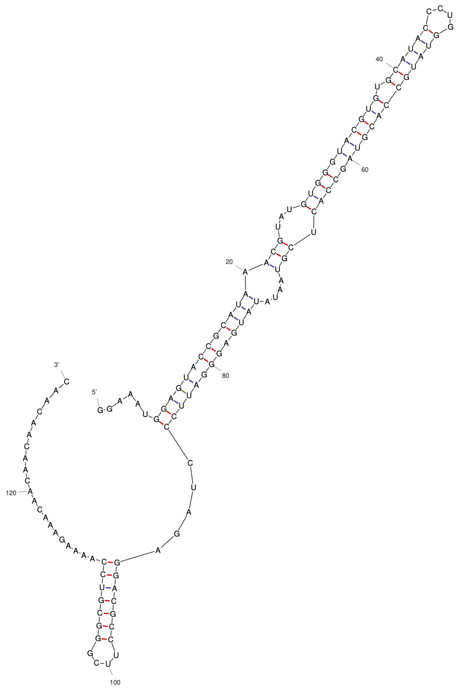
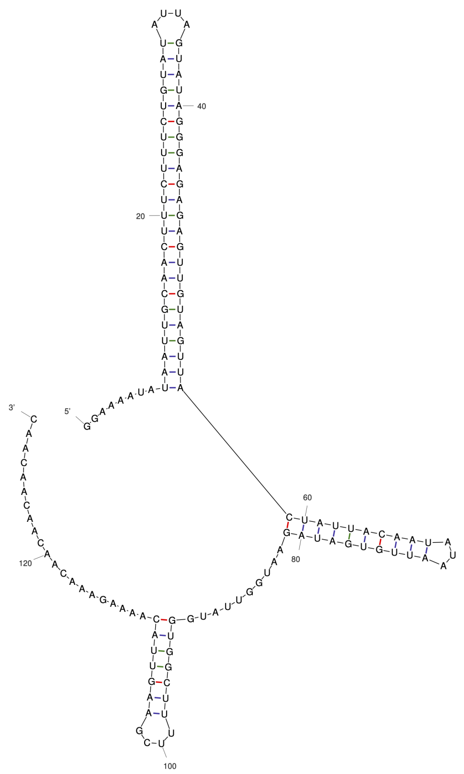
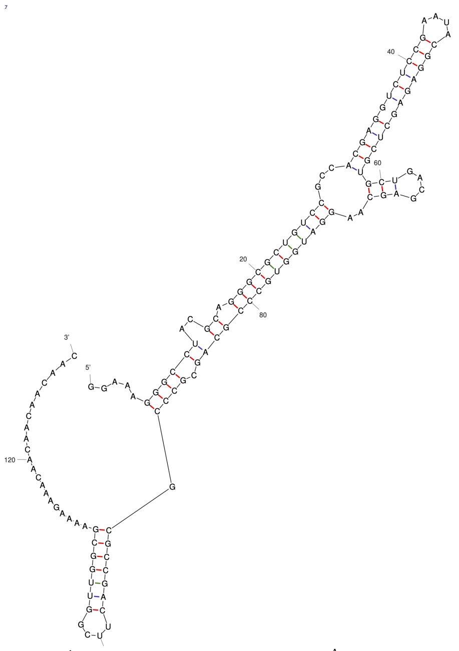
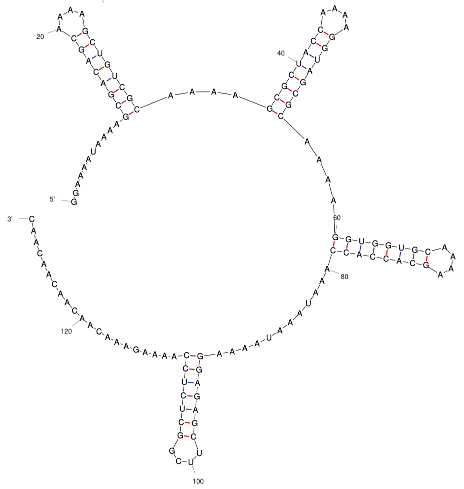
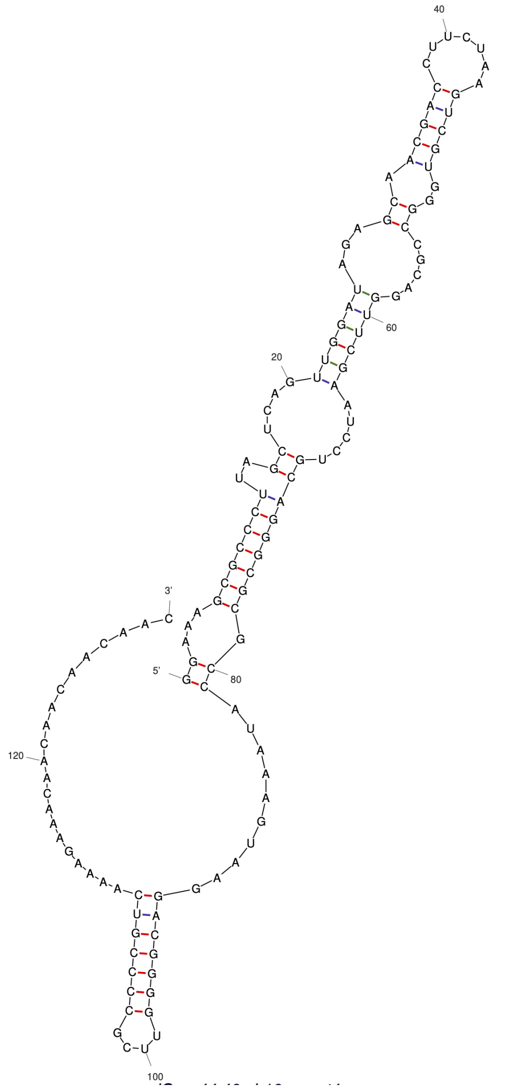
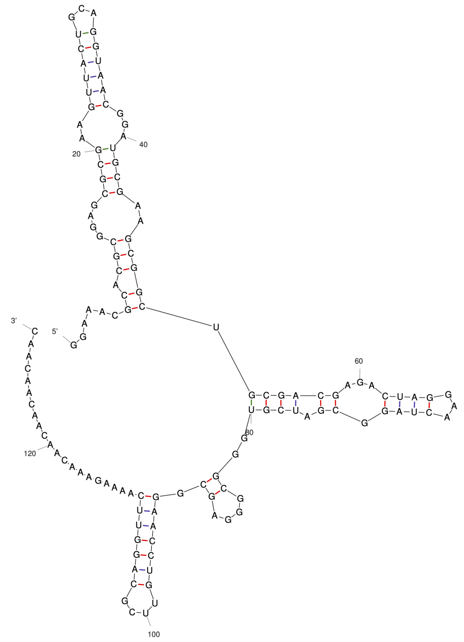
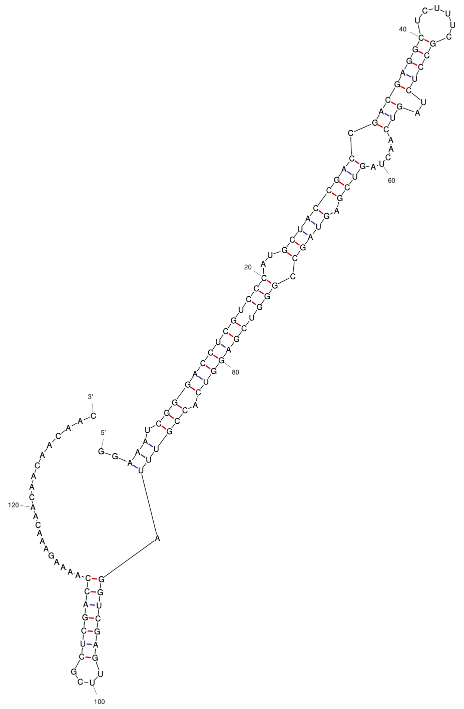
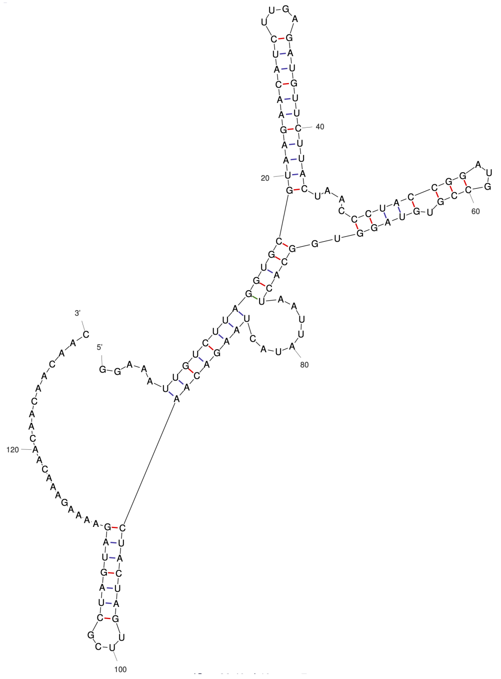
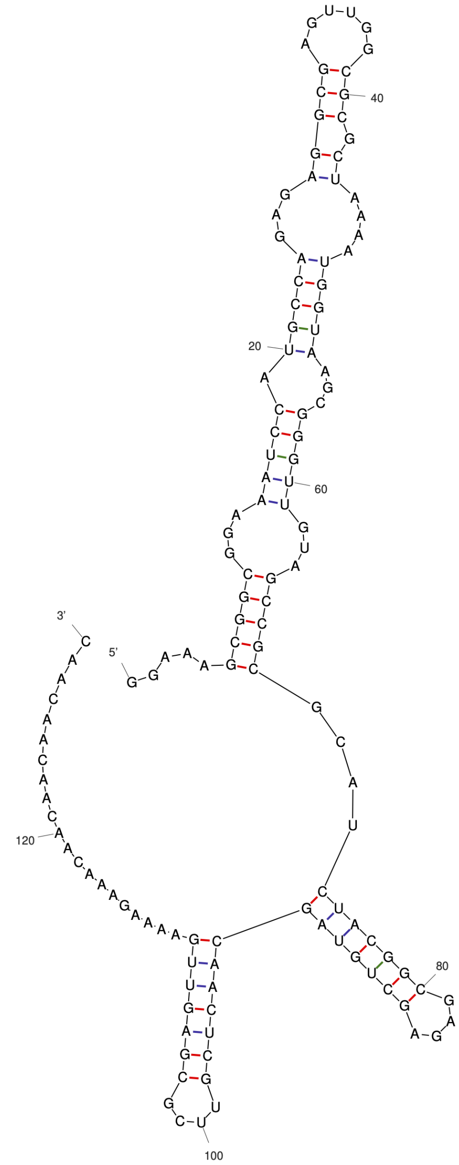
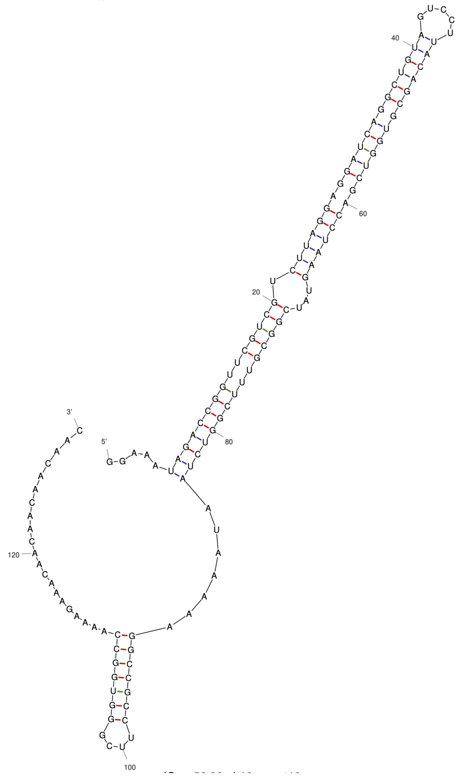

### Figure_S8.pdf

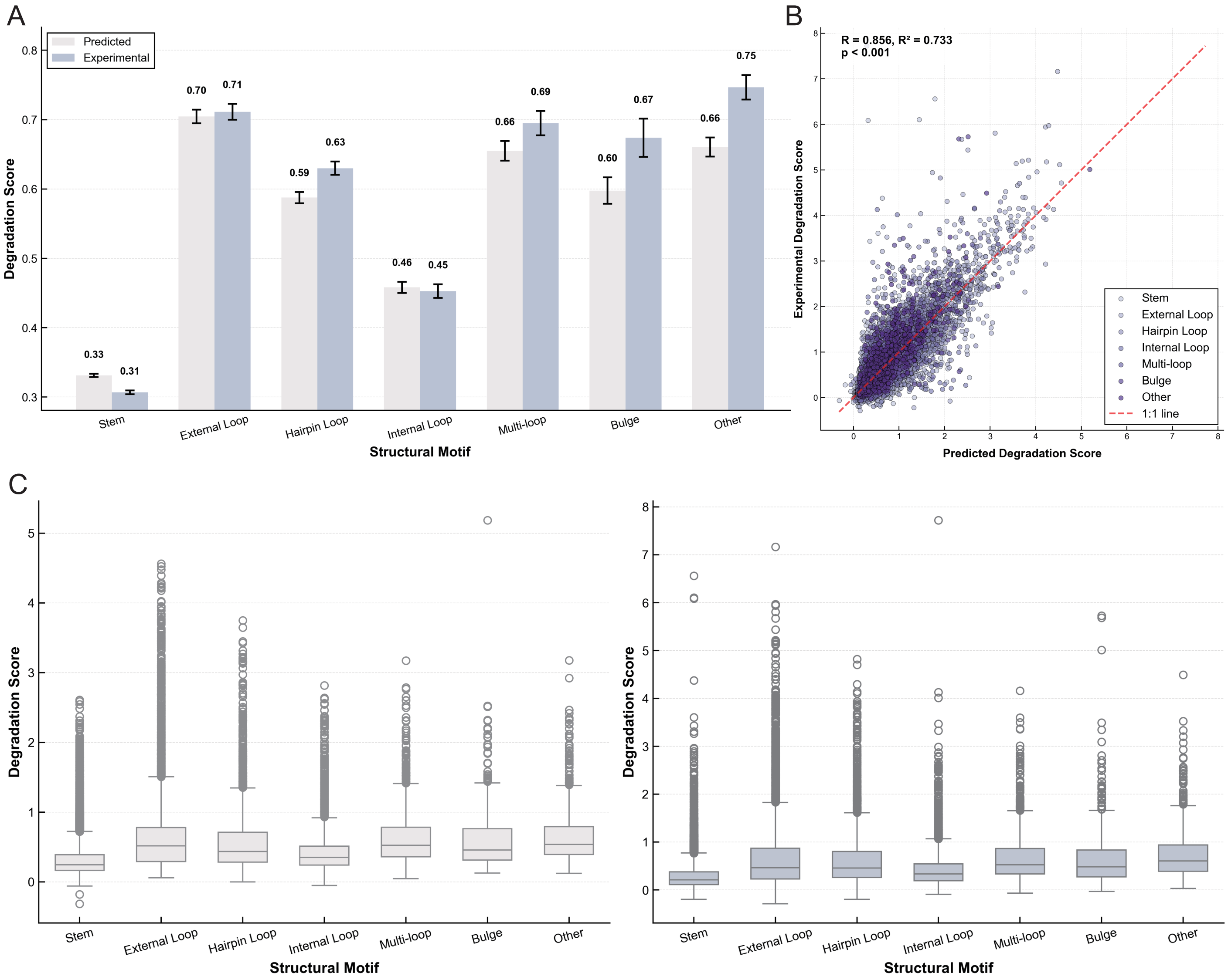

### Figure_S9.pdf

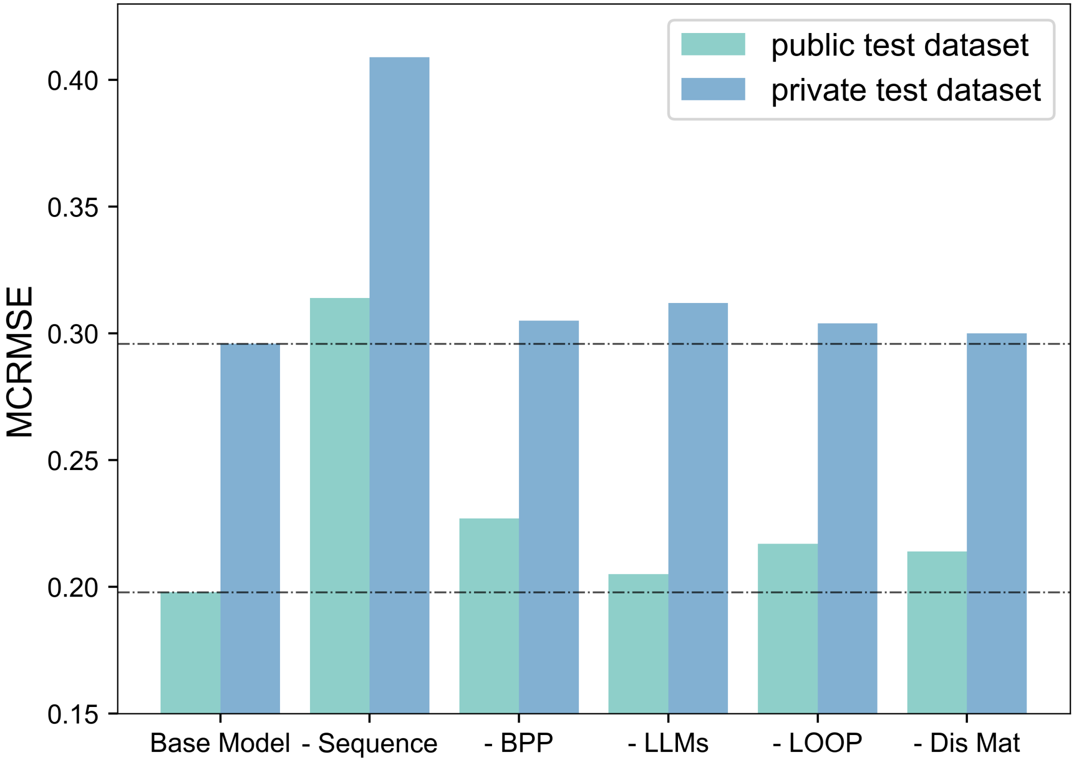
